# SCOTCH: isoform-level characterization of gene expression through long-read single-cell RNA sequencing

**DOI:** 10.1101/2024.04.29.590597

**Authors:** Zhuoran Xu, Hui-Qi Qu, Joe Chan, Shizhuo Mu, Charlly Kao, Hakon Hakonarson, Kai Wang

## Abstract

Recent advances in long-read single-cell transcriptome sequencing (lr-scRNA-Seq) enable full-length isoform profiling at single-cell resolution. We present SCOTCH (Single-Cell Omics for Transcriptome CHaracterization), an end-to-end, platform-independent pipeline for isoform characterization from lr-scRNA-Seq data, supporting Nanopore and PacBio sequencing as well as 10X Genomics and Parse Biosciences protocols. SCOTCH models isoforms as combinations of non-overlapping sub-exons and applies dynamic thresholding for robust isoform assignment while efficiently address ambiguous mapping issues. By refining sub-exon boundaries through integration of read coverage with existing annotations and applying an iterative clustering strategy to reconstruct novel transcripts, SCOTCH reliably recovers more true novel isoforms than existing splice-graph-based methods, with poly(A)-aware filtering further reducing false-positive structures. Extensive simulations demonstrate improved quantification of known isoforms and enhanced reconstruction of novel isoforms. Analyses of human blood and cerebral organoid datasets across multiple platforms further confirm SCOTCH’s ability to resolve cell-type-specific transcriptome profiles and uncover experimentally supported novel isoforms.

## Introduction

The development of single-cell RNA sequencing (scRNA-Seq) technology represents a transformative force in genomics, offering unprecedented insights into cellular heterogeneity and gene expression dynamics^1–3^. However, when paired with conventional short-read sequencing methods, scRNA-seq encounters limitations in fully capturing complex genomic regions and providing comprehensive view of transcriptomic diversity^4^. Although computational efforts such as SCALPEL^5^ have begun to characterize transcript diversity driven by alternative polyadenylation from 3’ tag-based scRNA-seq, these approaches remain constrained to 3’ transcript architecture and cannot resolve full-length isoforms or novel splicing events. In contrast, long-read sequencing technologies, such as those from PacBio and Oxford Nanopore Technologies, offer a significant advantage by spanning entire gene or transcript lengths ^6–9^. Consequently, the emergence of long-read scRNA-Seq (lr-scRNA-Seq) represents a significant leap forward, enabling isoform-level resolution of genetic analysis and offering a more nuanced view of the transcriptomic landscape within individual cells, which are key to understanding cellular function and disease mechanisms ^10–15^.

In recent years, various long-read single-cell technologies have been developed, including LR-Split-seq ^16^, Nanopore-specific adaptation of 10X Genomics Chromium^15^, FLASH-Seq ^17^, HIT-scISOseq^18^, MAS-ISO-seq^19^, among others, with each offering unique advantages. For instance, Parse Bioscience employs a split-pool combinatorial barcoding strategy, enabling the simultaneous sequencing of multiple samples. This approach is particularly advantageous for analyzing cell populations that are difficult to dissociate into single-cell suspensions, such as neurons, hepatocytes and developing cardiomyocytes. In addition, spatially resolved long-read transcriptomics has emerged as a powerful approach to capture isoform diversity within the spatial context of tissues^20, 21^. However, the emergence of long-read sequencing also introduces new challenges that require dedicated computational solutions. Although long-read data greatly reduce alignment ambiguity compared with short-read sequencing, some uncertainty still remains, motivating methods such as miniQuant^22^ to quantify read mapping uncertainty at isoform level. In addition, earlier Nanopore chemistries exhibited high error rates, complicating the process of identifying cell barcodes and unique molecular identifiers (UMIs), prompting the development of computational tools like LongCell ^23^ and FLAMES ^24^ to correct these inaccuracies and ensure the reliability of downstream data analysis. Many workflows therefore relied on paired short-read sequencing to guide barcode assignment ^25–27^. While with the introduction of R10 flowcells (v14 chemistry by Oxford Nanopore), per-base sequencing error rates have dropped to ~1% ^28, 29^, making previous error-focused computational methods less central nowadays. Instead, computational methods should shift their focus towards exploiting the unique advantages of long reads without short read guidance^30–34^: resolving transcript structures, modeling isoform usage, and more importantly, detecting novel isoforms accurately.

Although several computational tools have been developed for long-read transcriptomics, methods aimed at isoform quantification, including Bambu^35^, IsoQuant^36^, FLAMES^24^, and Isosceles^37^ still face notable challenges, particularly for novel isoform reconstruction and robust handling of ambiguous read alignments. These methods primarily rely on splice-graph reconstruction, which is highly sensitive to alignment noise and often generates fragmented transcript models. Subsequent filtering steps to manage these fragments, often based on static thresholds, may lead some methods to retain spurious novel isoforms and others to miss true ones, potentially limiting accurate novel isoform identification and annotation (as detailed in **Supplementary Notes**). In addition, most existing tools handle ambiguous read mappings by discarding uncertain reads or assigning them directly at the transcript level without resolving gene-level ambiguity first, contributing to substantial read loss and reduced accuracy in isoform quantification. Furthermore, existing tools each face distinct challenges in the single-cell context. Bambu^35^ and IsoQuant^36^, originally designed for bulk RNA-seq data, can be adapted for single-cell applications but requires extensive manual preprocessing steps such as read de-multiplexing and barcode handling, which complicate their use in automated workflow. FLAMES^24^, while specifically designed for lr-scRNA-Seq data, still depends on short-read data for barcode identification, limiting its use as a long-read-only tool, despite its capabilities in isoform discovery and splicing analysis. Isosceles^37^, another tool designed for transcript discovery and quantification in lr-scRNA-Seq, does not accommodate the diverse read structures from different single-cell platforms. Many of these existing methods use statistical models that compare exon inclusion levels across cell populations, limiting their ability to directly assess isoform usage differences at the transcript level—an essential aspect for fully understanding transcriptome complexity. In this context, the development of new computational methods that can fully exploit the strengths of lr-scRNA-Seq, while overcoming these limitations, is critically needed.

In this study, we generated a comprehensive set of benchmarking datasets using both Illumina short-read and Oxford Nanopore long-read sequencing, encompassing different flow cells versions (R9 and R10) and single cell preparation protocols from 10X Genomics and Parse Biosciences. Our examination of lr-scRNA-seq data has shown that recent technological advancements have eliminated the need for parallel short-read sequencing, rendering previous concerns about high error rates less relevant. To address the unique challenges of lr-scRNA-seq, we introduce Single-Cell Omics for Transcriptome CHaracterization (SCOTCH), a suite of computational pipeline and statistical framework specifically designed for lr-scRNA-seq transcriptome analysis. SCOTCH is compatible with single-cell libraries from 10X Genomics and Parse Biosciences, as well as sequencing platforms from Nanopore and PacBio. It excels in detecting transcript usage differences between cell populations, identifying isoform switching events, and discovering novel isoforms. By refining exon boundaries using read-coverage profiles and applying Louvain clustering to group reads into coherent candidate transcript structures followed by iterative realignment, SCOTCH mitigates the fragmentation issue commonly observed in splice-graph–based methods, enabling improved novel isoform recovery with reduced transcript model redundancy, while accounting for poly(A) stretch-related artifacts. SCOTCH also employs a read-specific dynamic thresholding strategy that integrates read mapping scores to resolve multi-mapping reads from overlapping genes and chimeric transcripts, thereby enhancing robustness to sequencing and alignment noise and ensuring accurate isoform quantification with minimal read loss. Through extensive simulation studies, we demonstrate SCOTCH’s superior performance in isoform annotation and quantification for both known and novel isoforms, and its ability to handle complex mapping challenges, such as overlapping genes and chimeric reads. By applying SCOTCH to human PBMC (peripheral blood mononuclear cell) and cerebral organoid datasets, we demonstrate SCOTCH’s capability to reveal novel biological insights from lr-scRNA-seq data, providing transcript-level resolution that short-read data cannot achieve. Experimental validations, including PCR and qPCR assays, further confirmed the accuracy of SCOTCH in isoform quantification and novel isoform identification. Our study highlights lr-scRNA-seq’s potential to deepen our understanding of genome regulation and cellular diversity at single-cell resolution. Additionally, the datasets generated from five distinct technical approaches provide a valuable resource for further development of computational methods.

## Results

### SCOTCH workflow and experimental design

In the current study, we introduce SCOTCH, a suite of computational pipeline and statistical framework specifically developed for analyzing lr-scRNA-seq data (**Figure 1)**. SCOTCH preprocessing pipeline supports libraries generated by both 10X Genomics and Parse Biosciences paired with Nanopore or PacBio long read sequencing (**Figure 1a**), with other library generation protocols in the works. For 10X Genomics libraries, SCOTCH detects polyA/T tails connected to UMI and cell barcodes, and account for possible read truncations at the 5’ end. For the Parse libraries, SCOTCH similarly accounts for possible 5’ read truncations if polyA/T tails can be detected in the long reads; otherwise, it considers read truncations at both 3’ and 5’ ends when hexamer primers might be used.

**Figure 1.**
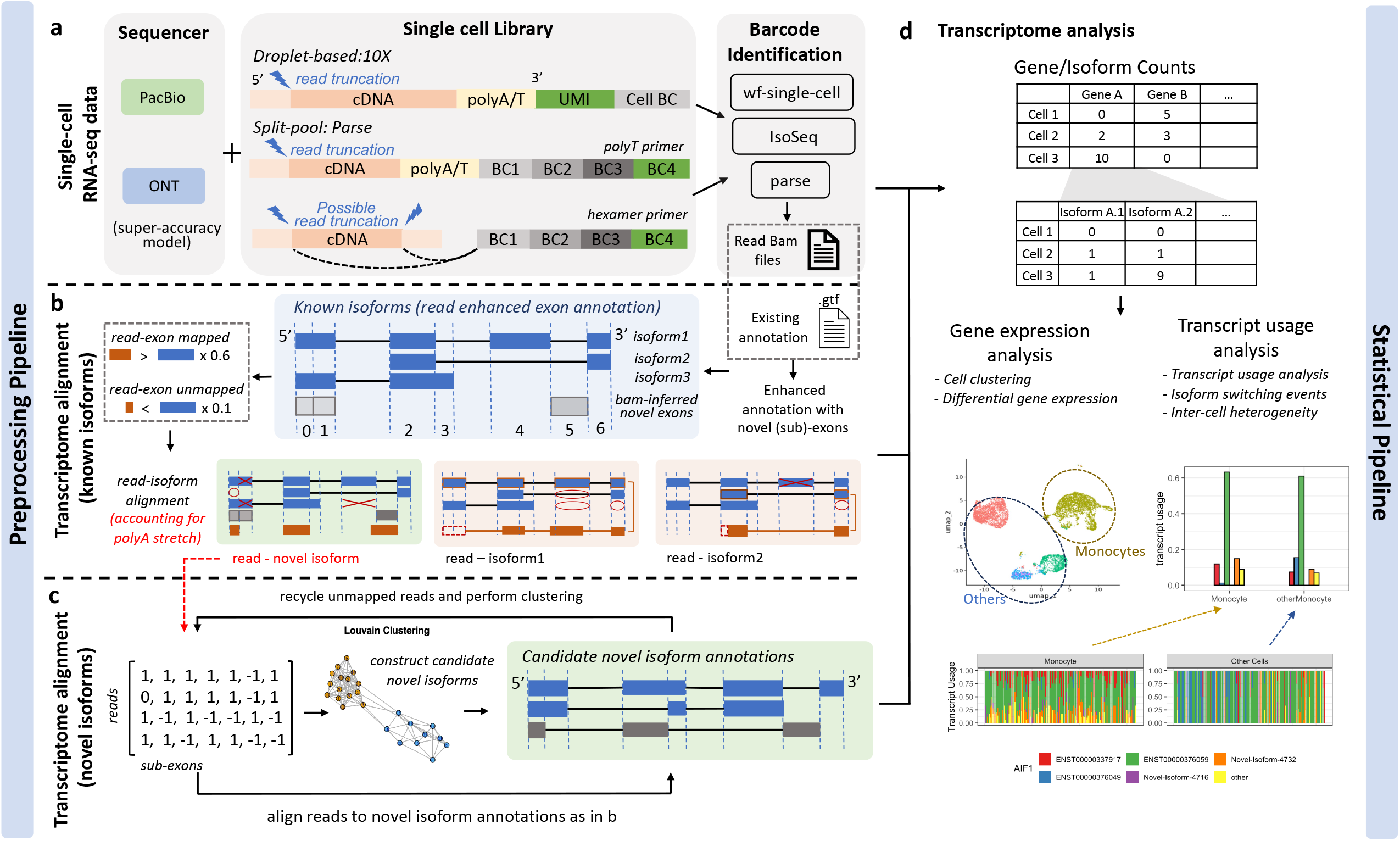
Workflow of the SCOTCH pipeline for long-read scRNA-seq analysis. (a) For either the 10X Genomics or Parse Biosciences single-cell library construction method, the cell barcodes and UMIs are extracted using vendor-supplied protocols. SCOTCH accounts for potential read truncations depending on primers used. (b) SCOTCH dissects known isoform annotations into non-overlapping exons and integrates read coverage information from BAM files to refine these annotations by identifying novel candidate exons or splitting existing exons into sub-exons. SCOTCH then maps reads to each sub-exon, applying a threshold where reads are considered mapped if it covers more than 60% of the exon’s length, and considered unmapped if it covers less than 10%. Using these read-exon mapping calls, reads are then aligned to known isoforms through a compatible matrix. (c) Reads that cannot be mapped to any known isoforms are iteratively clustered to generate candidate novel isoform annotations at the pseudo-bulk level based on read-exon matching profiles followed by read-isoform remapping. (d) lr-scRNA-seq data generated using 10X Genomics or parse libraries can be preprocessed by vendor-supplied tools, such as the wf-single-cell or Parse pipelines for barcode identification and then processed by SCOTCH to generate count matrices at both the gene and transcript levels. The gene-level count matrix facilitates conventional gene expression analysis, while the isoform-level count matrix can be used for transcript usage analysis through SCOTCH’s statistical pipeline.

SCOTCH takes vendor-tagged BAM files (e.g., wf-single-cell for 10X-ONT, Iso-seq for 10X-PacBio, and Parse pipeline for Parse-ONT data) as input to align reads to known and candidate novel isoforms, each presented as combinations of non-overlapping sub-exons defined from annotation file (**Figure 1b-c**). Specifically, SCOTCH encodes reads by the presence or absence of each sub-exon, applying a dynamic thresholding scheme based on read-exon mapping percentages and accounting for poly(A) stretch-related artifacts to accommodate varying sequencing and alignment noise across transcriptome. Read-isoform mappings are discarded if (1) the isoform annotation includes any sub-exons that the read skips or (2) the isoform annotation fails to encompass all exons covered by the read. For sub-exon annotations, SCOTCH supports three modes: annotation-only (using only existing annotation files), annotation-free (relying solely on read coverage information), and enhanced-annotation (the default mode, combining both existing annotations and read coverage to refine annotations with novel sub-exon boundaries, see **Methods** for more details). This flexibility increases sensitivity for discovering novel isoforms, particularly those involving novel exons, including intron retentions and 5’/3’ alternative splicing events. After initial read alignment to known isoforms (**Figure 1b**), SCOTCH builds a read-read similarity graph for the remaining unmapped reads based on read-exon matching profiles and applies the Louvain clustering method^38^ to identify coherent read groups, from which candidate novel isoform structures are inferred at the pseudo-bulk level. SCOTCH then realign these reads to the resulting candidate models and leverage multi-mapping evidence to consolidate redundant or truncation-derived structures, preventing inflation of the novel isoform set (**Figure 1c** and **Supplementary Notes**). This iterative refinement provides a more coherent novel isoform annotation than splice-graph–based approaches, while remaining resilient to sequencing and alignment noises. Notably, A read may map to multiple transcripts of a gene or, in more complex cases, ambiguously map to several overlapping genes or even nonoverlapping genes, as seen with chimeric reads. To resolve these ambiguities, SCOTCH adopts read mapping scores (see **Methods**) to prioritizes the most likely transcript or gene assignment.

After preprocessing lr-scRNA-seq data, we aim to perform analysis with transcript-level resolution to unveil alterations obscured by conventional differential gene expression analysis. Differential transcript usage (DTU), which is defined as variations in relative abundance of transcripts of the same gene across different conditions or cell types, explain changes in phenotype between cell types, tissues, or disease cohorts ^39–41^. The SCOTCH statistical pipeline, which is inspired by LIQA ^42^ for truncation handling and LongCell ^23^ for exon-inclusion analysis, can be used to identify DTU on both the gene and transcript levels (**Figure 1d, Methods**). For each gene, SCOTCH estimates the average transcript usage of a cell population by fitting a Dirichlet-multinomial distribution, which captures inter-cell variations quantified by the over-dispersion parameter ϕ. This parameter is mean invariant, where small values indicate cells with similar isoform co-expression patterns, and large values suggest a more exclusive expression mode, where isoform usage patterns are more variable across cells. Employing likelihood ratio tests, we assess whether transcript usage differ between two cell populations at the gene level, and we examine whether a specific transcript is differentially utilized at the transcript level. One special case of transcript usage alteration is isoform switching, here defined as changes of the dominant isoform between two cell populations (we note that it may also be denoted as a synonym for DTU in some publications). SCOTCH is designed to pinpoint isoform switching events, whose effect size is measured by the absolute sum of differences in the dominant isoform proportions between the two cell populations.

To benchmark our methods and assess how SCOTCH aids in lr-scRNA-seq analysis for understanding transcriptome complexity, we first simulated lr-scRNA-seq data for two distinct cell populations with ground truth. Additionally, we generated a substantial amount of evaluation data for two human peripheral blood mononuclear cells (PBMC) samples using both 10X Genomics and Parse libraries, Illumina and nanopore sequencings (**Table 1**). Throughout the study, we first demonstrated that the availability of R10 flowcells facilitates isoform-level analysis without paired short reads or complex computational methods previously required to handle high sequencing error rates. This is shown through several key comparisons, including short-read versus long-read sequencing, R9 versus R10 flowcells sequencing technologies, and the 10X Genomics versus the Parse Bioscience library preparation system. Further into the study, we compared SCOTCH’s performance with several benchmark methods and analyzed lr-scRNA-seq data from PBMCs and human cerebral organoids^32^ to showcase the effectiveness of SCOTCH’s statistical framework in analyzing DTU, detecting isoform switching events, identifying novel isoforms, and assessing inter-cell heterogeneity.

**Table 1.** Summary of human PBMC benchmarking datasets generated in the current study.

| Sample Name | Sequencing Platform | Flowcell | Single Cell Library | Number of Reads<br>Pass Filter | Mean Q-scores |
| --- | --- | --- | --- | --- | --- |
| Sample 7 | Illumina |  | 10x Genomics 3' v3 | 313,271,568 | 35.82 |
| Sample 7 | Illumina |  | Parse Biosciences | 776,207,860* | - |
| Sample 7 | Nanopore | R9 | 10x Genomics 3' v3 | 123,276,129 | 10.23 |
| Sample 7 | Nanopore | R10 | 10x Genomics 3' v3 | 138,389,424 | 11.19 |
| Sample 7 | Nanopore | R10 | Parse Biosciences | 230,564,859* | 12.66* |
| Sample 8 | Illumina |  | 10x Genomics 3' v3 | 308,786,488 | - |
| Sample 8 | Illumina |  | Parse Biosciences | 776,207,860* | 35.71 |
| Sample 8 | Nanopore | R9 | 10x Genomics 3' v3 | 132,392,128 | 10.33 |
| Sample 8 | Nanopore | R10 | 10x Genomics 3' v3 | 172,081,920 | 11.45 |
| Sample 8 | Nanopore | R10 | Parse Biosciences | 230,564,859* | 12.66* |

\*: The statistics for Parse Biosciences samples represent those across all four samples.

### R10 flowcells facilitate isoform-level analysis with improved sequencing quality

When preprocessing lr-scRNA-seq data with library constructed on the 10X Genomics platform, a threshold of edit distance (ED) is set for UMI clustering and cell barcode resolution. This ED threshold determines the stringency of the error correction steps for UMIs and cell barcodes, directly impacting the precision of read allocation to particular cells and the effectiveness of molecule de-multiplexing. The conventional wisdom is that R9 flowcells need an ED of 2 for accurate read-cell mapping ^43^, owing to their higher sequencing errors compared to short-read sequencing. However, the newer R10 flowcells display improved sequencing quality, particularly in homopolymers regions ^44^, where they have lower deletion rates in thymine and adenine sequences than R9, albeit with increased mismatches ^45^. The R10’s ability to match Illumina’s sequencing quality standards yields flexibility in choosing between ED1 and ED2, allowing for more efficient and accurate read mappings to the corresponding cells.

To empirically substantiate the technical advancements offered by R10 flowcells that facilitate isoform-level analysis, we systematically examined and compared sequencing statistics across different combinations of single-cell libraries and sequencing technologies (10X + Illumina, Parse + Illumina, 10X + Nanopore_R9, 10X + Nanopore_R10, Parse + Nanopore_R10) that are preprocessed by vendor-supplied computational pipelines (**Table 1, Table S1, Figure 2**). We observed higher number of reads passing the filter with higher mean Q-scores of R10 flowcells compared of those of R9 flowcells. Regardless of whether an ED of 1 or 2 is applied, R10 flowcells align more closely with short-read sequencing in terms of valid barcode numbers (**Figure S9-S11**), median numbers of genes and UMIs per cell, as well as the total counts of cells, genes, and transcripts (**Table S1, Figure 2a**). R10 flowcells also show higher proportions of reads categorized as full length, total-tagged, gene-tagged, and transcript-tagged (**Figure 2b**), indicating higher sequencing quality. When applying an ED of 2, both R9 and R10 flowcells display an increase in sequencing saturation and elevated median gene, transcript, and UMI counts per cell compared to using ED of 1. This is because a higher ED threshold leads to more lenient clustering, thereby enabling the recognition of more unique transcripts. Notably, the increase seen with ED2 relative to ED1 is less pronounced for R10 than R9, and both ED1 and ED2 exhibit comparable performance in terms of the total numbers of cells, genes, and transcripts for R10 (**Figure 2a, Figure S1**), suggesting that the improved sequencing accuracy of R10 flowcells reduces the need for a higher ED for reliable transcript identification. Additionally, R10 consistently outperforms R9 in sequencing saturation, and median numbers of genes, transcripts, and UMI counts per cell across both ED settings, affirming its ability to capture a more comprehensive transcriptome profile even at a stricter ED threshold.

**Figure 2.**
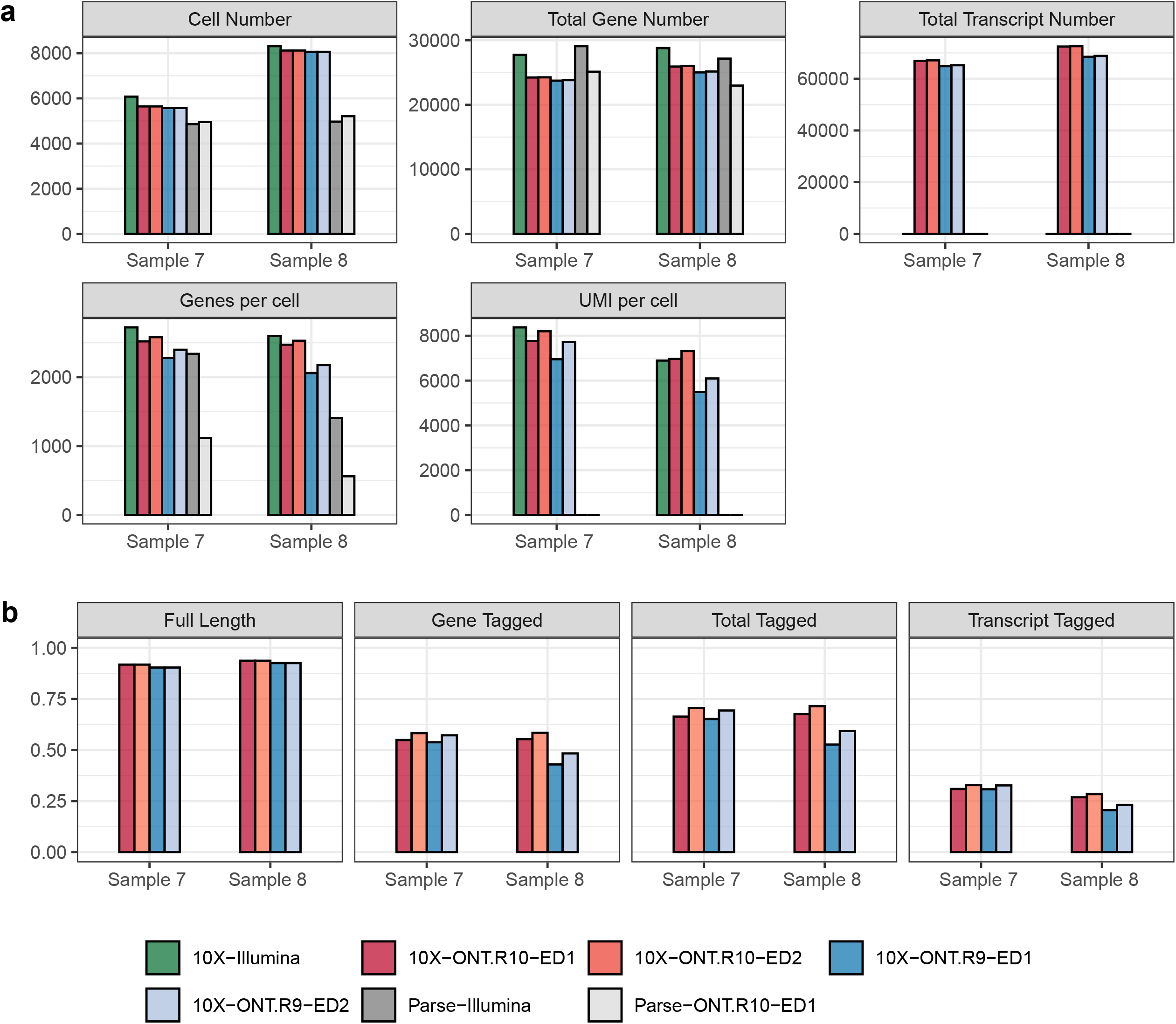
Influence of edit distance (ED) criteria on final results of five single-cell library + sequencing platforms generated by vendor computational pipeline: 10X + Illumina, Parse + Illumina, 10X + Nanopore_R9, 10X + Nanopore_R10, Parse + Nanopore R10. (a) Cell number, total gene number, total transcript number, median number of genes per cell, and median number of UMI per cell that are identified based on different flowcell versions and different ED thresholds (1 or 2). (b) The fraction of reads that are classified as full-length, gene-tagged, total-tagged, and transcript-tagged in two samples sequenced by both R9 and R10 flowcells on the Nanopore platform. Source data are provided as a Source Data file.

### SCOTCH accurately characterizes and quantifies transcripts in simulation studies

To comprehensively evaluate SCOTCH performance in characterizing and quantifying transcripts, as well as supporting downstream transcriptome analysis, we simulated ground-truth nanopore scRNA-seq data for two distinct cell populations using a modified version of lrgasp-simulation specifically adapted for single-cell data (see **Methods**). Specifically, we simulated reads from 1080 expressed genes on chromosome 6, with each having at least two gene isoforms based on GENCODE gene annotation^46^. To examine the ability of SCOTCH in the discovery and annotation of novel isoforms, we randomly removed 2466 (30%) isoform structures, designating them as ground-truth novel isoforms, while the remaining 5762 (70%) isoforms in all 1080 genes were used as the existing annotation file for all tools. To assess whether SCOTCH’s gene- and transcript-level count matrices preserved underlying differential transcript usage signals relevant to disease studies, we set the transcript compositions of 540 (50%) genes to differ between two cell populations, with the remaining 540 genes maintaining the same transcript compositions across both groups. In total, we simulated 15,381,985 reads for two cell populations, each having 1000 cells. We compared performance of SCOTCH (default enhanced-annotation mode), SCOTCH.ao (annotation-only mode), and several other methods including IsoQuant, Bambu, FLAMES (modified by us to process single-cell data), and Isosceles (with both strict and loose modes).

We first examined the composition of mapped reads for each method. As shown in **Figure 3a**, most competing methods display a significant proportion of ambiguously mapped reads, in some cases even surpassing the number of unique mappings and exhibit a notable number of missing reads. This is a known challenge in long-read single-cell data, and it has influences on downstream data analysis since a large proportion of reads cannot be assigned to transcripts. In contrast, SCOTCH and SCOTCH.ao achieve the highest number of uniquely mapped reads with minimal ambiguity, showcasing a unique advantage of handling ambiguous mapping. We then compared read-isoform mapping accuracy for both known and novel isoforms. **Figure 3b** shows that SCOTCH, in enhanced-annotation mode, achieves accuracy of 0.876, comparable to IsoQuant (0.878). The annotation-only mode (SCOTCH.ao) also performs well with an accuracy of 0.793, significantly outperforming other tools such as FLAMES (0.461) and Bambu (0.163). The accuracy difference between SCOTCH and SCOTCH.ao likely stems from SCOTCH’s ability to refine sub-exon annotations using read coverage information, enhancing its ability in novel isoform detection. To further explore this, we treated read-isoform mappings as a classification task to predict whether a read originates from a novel isoform. As seen in **Figure 3c**, SCOTCH accurately detects novel isoforms, with an F1 score (0.815) slightly lower than IsoQuant (0.853). SCOTCH and SCOTCH.ao exhibit similarly high precision (SCOTCH: 0.984, SCOTCH.ao: 0.981), but SCOTCH demonstrates significantly higher recall (0.695) than SCOTCH.ao (0.413), indicating its enhanced sensitivity in detecting novel isoforms. In comparison, Bambu and FLAMES show much lower precision and recall. Isosceles does not report read-isoform mappings and was thus excluded from these comparisons. We then evaluated the performance of various tools in annotating novel isoforms. We excluded ground-truth novel isoforms with bulk-level counts less than 20, as such low-abundance transcripts are difficult to reliably detect and are rarely considered in practical analyses. To ensure a fair comparison, we uniformly removed predicted novel isoforms with estimated expression levels below 20 across all tools prior to performance assessment. After filtering, 2391 truth novel isoforms were retained. To assess annotation accuracy, we used gffcompare^47^ to compare GTF files generated by each tool with ground truth and calculated **precision** and **recall** (**Figure 3e**; see **Supplementary Notes**). Briefly, precision (prediction-centric) quantifies the fraction of predicted novel isoforms that confidently match true novel isoforms, whereas recall (truth-centric) measures the fraction of true novel isoforms recovered, with both metrics based on high-confidence isoform-level matches. We also report transcript model redundancy, defined as the average number of predicted transcript models corresponding to each true novel isoform, as excessive transcript model fragmentation can inflate transcript-level variability estimates and obscure biologically meaningful signals even when overall annotation accuracy improves. As shown in **Figure 3d**, IsoQuant (698) and Bambu (404) recovered only a limited number of the ground-truth novel isoforms, resulting in low recall (**Figure 3j**; IsoQuant: 0.257, Bambu: 0.156). However, novel transcript models they constructed were highly reliable, achieving high precision (**Figure 3j**; IsoQuant: 0.907; Bambu: 0.978) with minimal redundancy, producing approximately one predicted transcript model per true novel isoform (**Figure 3e**). On the other hand, Isosceles.strict (1814), Isosceles.loose (2353), and FLAMES (1822) reported numbers of novel isoforms closer to the ground truth (**Figure 3d**). Nevertheless, this apparent numerical proximity did not translate into a substantial improvement in true isoform recovery. Isosceles.strict (recall = 0.311) and Isosceles.loose (recall = 0.392) achieved moderately higher recall than IsoQuant and Bambu, whereas FLAMES recovered an even smaller fraction of true novel isoforms (recall = 0.220). These changes in recall came at the cost of substantial redundancy (**Figure 3e**) and increased false positive predictions, with nearly two transcript models corresponding to each true novel isoform on average and approximately 10% of predicted models failed to match any ground-truth novel isoforms (precision: Isosceles.strict: 0.770; Isosceles.loose: 0.781; FLAMES: 0.531). In contrast, SCOTCH identified 1794 novel isoforms while recovering a markedly larger fraction of true novel transcripts (recall = 0.561), more than doubling the recall of IsoQuant and exceeding all other methods. Importantly, this improvement in sensitivity did not compromise annotation quality. SCOTCH maintained low redundancy (1.13 predicted transcript models per true novel isoform; **Figure 3e**), and a low false positive rate, with fewer than 5% of predicted novel isoforms failing to match ground truth, resulting in high precision of 0.882. Consequently, SCOTCH achieved the highest overall F1 score (0.686) outperforming all other methods (SCOTCH.ao: 0.450, IsoQuant: 0.400, FLAMES: 0.311, bambu: 0.268, Isosceles.strict: 0.443, Isosceles.loose: 0.522).

**Figure 3.**
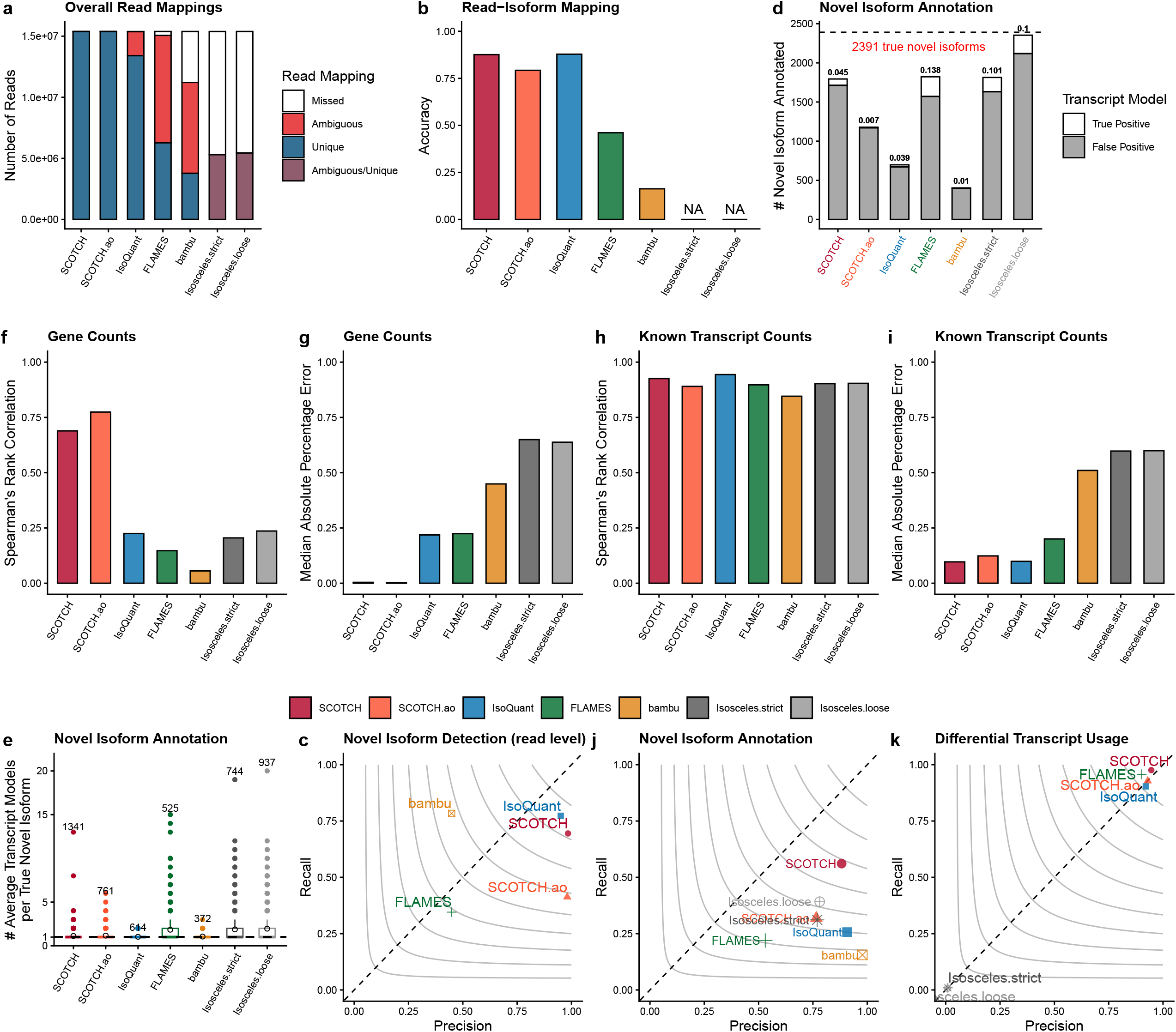
Simulation study to evaluation SCOTCH performance compared with benchmark methods. (a) Bar plots showing number of reads that mapped to one isoform uniquely (unique mapping), multiple isoforms ambiguously (ambiguous mapping), and missed by tools. (b) Accuracy of read-isoform mappings including known and novel transcripts. Since Isosceles does not output read-isoform mapping information, NA was annotated. (c) Precision and recall for classifying reads to known or novel isoforms. (d) Number of novel isoforms identified by different methods. The dashed horizontal line indicates the 2,391 ground-truth novel isoforms. Predicted transcript model that can be matched to a true novel isoform by GffCompare is true positive, otherwise false positive. Numbers above the bars indicate the false positive rate (FPR). (e) Boxplots showing distribution of matched number of predicted transcript models per true novel isoform. The dashed line indicates the ideal one-to-one correspondence. Boxplots show the median (center line), interquartile range (box), and values within 1.5 × IQR (whiskers), with points beyond this range being outliers. Black triangle markers indicate the mean values. Numbers above each boxplot denote the sample size of each distribution, representing the number of true novel isoform being recovered by high-quality transcript models. (f) Spearman’s rank correlation of gene expressions between ground-truth values and tools’ output for 1080 genes on the bulk level. (g) Median absolute percentage error of gene expressions between ground-truth values and tools’ output for 1080 genes on the bulk level. (h) Spearman’s rank correlation of transcript expressions between ground-truth values and tools’ output for 5762 known transcripts on the bulk level. (i) Median absolute percentage error of transcript expressions between ground-truth values and tools’ output for 5762 transcripts on the bulk level. (j) Precision and recall for annotating novel isoforms. (k) Precision and recall for identifying DTU genes. DTU genes were identified using two-sided likelihood ratio tests with Holm-adjusted P values < 0.01 considered significant. (d-e, j) contour lines represent F1 scores. Source data are provided as a Source Data file.

After evaluating transcript characterization, we shifted our focus to quantification, comparing the correlations between gene (**Figure 3f-g**) and known transcript counts (**Figure 3h-i**) generated by each tool and ground truths. SCOTCH and IsoQuant exhibit similarly high correlations with ground truth (SCOTCH: 0.926, IsoQuant: 0.944), and low MAPE values (SCOTCH: 0.097, IsoQuant: 0.099) for known transcripts. While other tools perform worse in transcript quantification, the differences are not as pronounced. However, for gene-level quantifications, SCOTCH and SCOTCH.ao significantly outperform other tools, achieving correlation coefficients of 0.689 and 0.774, respectively, both with remarkably low MAPE values (SCOTCH: 0.004, SCOTCH.ao 0.003). In comparison, Isosceles.loose and IsoQuant only achieved correlations of 0.236 and 0.225, with MAPE being 0.638 and 0.218, with other tools performing even worse. This superior performance can be attributed to SCOTCH’s ability to resolve ambiguous mappings more effectively, as seen in **Figure 3a**, where SCOTCH showed a higher proportion of uniquely mapped reads with minimal ambiguity. Although gene-level quantification is not the primary motivation for using long-read sequencing, this result highlights that SCOTCH achieves accurate quantification at both gene and isoform levels, making it unnecessary to rely on additional short-read data for improved accuracy. We finally assessed how each tool aids in transcriptome analysis by preserving the underlying differential transcript usage (DTU) signals (**Figure 3k**). SCOTCH, SCOTCH.ao, FLAMES and IsoQuant closely align with the ground truth in terms of precision and recall, with SCOTCH outperforming the others. However, Isosceles.loose and Isosceles.strict performs the worst, with near-zero precision and recall, likely due to the vast number of missed and ambiguous reads (**Figure 3a**). These results demonstrate SCOTCH’s effectiveness in accurately capturing DTU signals, further supporting its robustness in transcript-level analysis.

### Consistency and reproducibility between different sequencing technologies, single-cell libraries, and computational pipelines

After the simulation studies, we applied SCOTCH, SCOTCH.ao, IsoQuant, and the wf-single-cell pipeline to two human PBMC samples processed with different single-cell libraries and sequencing protocols. Due to Isosceles’s high rate of missed reads, FLAMES’ reliance on short-read barcoding, Bambu’s focus on bulk sequencing, and Parse’s inability to generate transcript-level matrices (as of November 2024), these tools were excluded from this analysis. We first compared cell type clusters on the gene level across short-read and long-read scRNA-seq, 10X Genomics and Parse single-cell libraries, R9 and R10 flowcells, and with Edit Distance of 1 and 2, as well as different computational pipelines using PBMC samples (**Figure 4a-c**). The cell type clusters were generally consistent in UMAP visualizations and cell type proportions across different platforms and analytical methods, with monocytes being the most prevalent cell type, followed by B cells. These patterns were reproducible across two samples, underscoring the reliability of cell type identification across diverse sequencing technologies, library preparations, and analytical approaches on the gene level. On the transcript level, we assessed the number of DTU genes across two samples identified by different computational methods, utilizing data generated by 10X Genomics library with nanopore R10 flowcells (ED of 1). As shown in **Table 2**, the SCOTCH and SCOTCH.ao consistently identified a greater number of DTU genes across different cell types than the wf-single-cell pipeline for most cell types, with IsoQuant detecting significantly fewer DTU genes. Moreover, the results from the second sample confirmed the superior reproducibility of SCOTCH and SCOTCH.ao, with both methods maintaining a higher percentage of reproducible DTU genes for most cell types across samples compared to other tools (**Figure 4d**). Owing to the low sequencing depth of the Parse library, we proceeded to determine whether the Parse platform enables the identification of key DTU genes that were also detected using the 10X Genomics platform. We found that over half of DTU genes detected in B, monocytes, and NK cells by Parse-SCOTCH were validated by 10X-SCOTCH (**Figure S3**), suggesting strong consensus between both single-cell library construction methods.

**Table 2.**
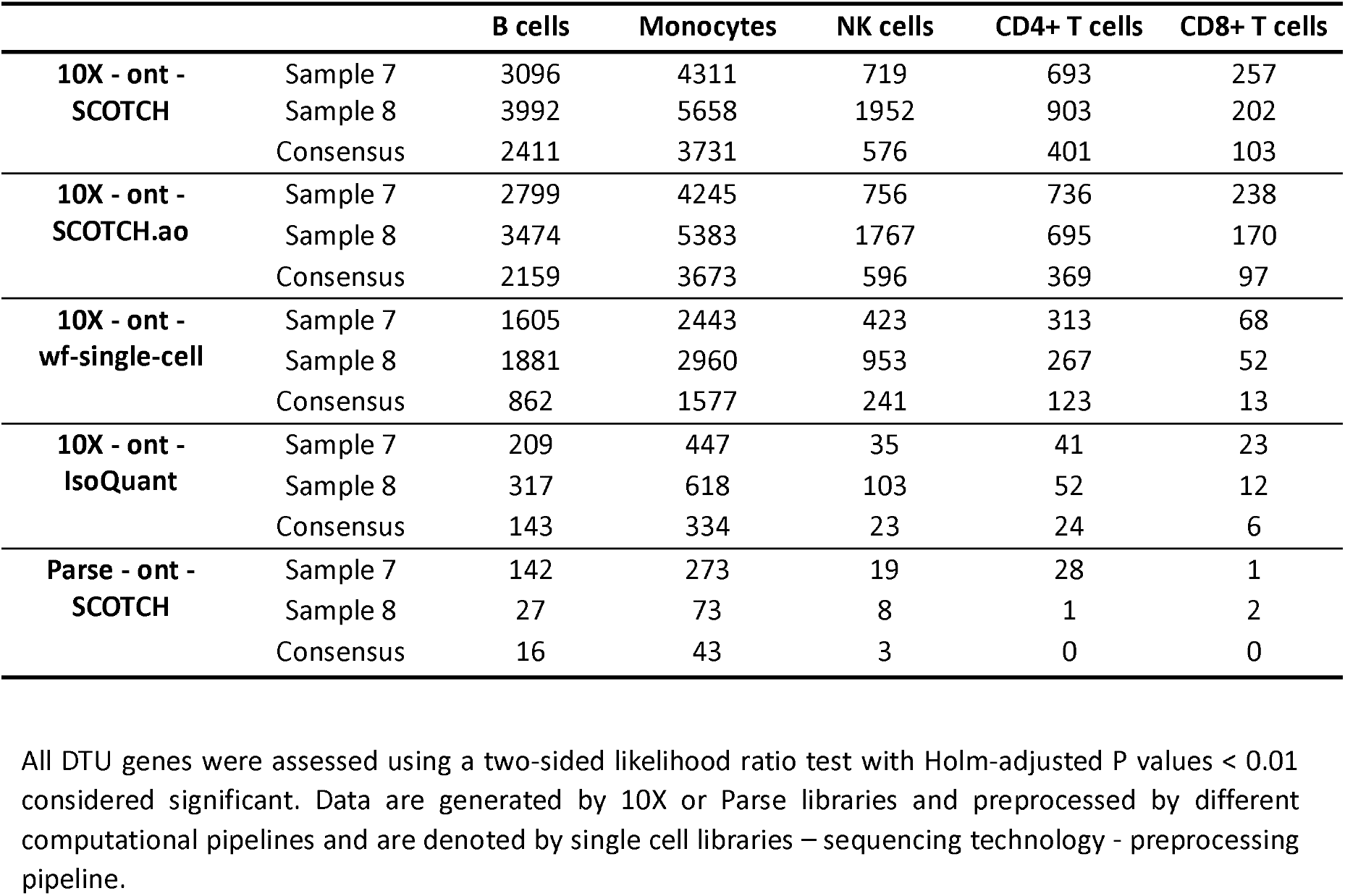
Number of genes with significant differential transcript usage (DTU) identified by different computational tools.

**Figure 4.**
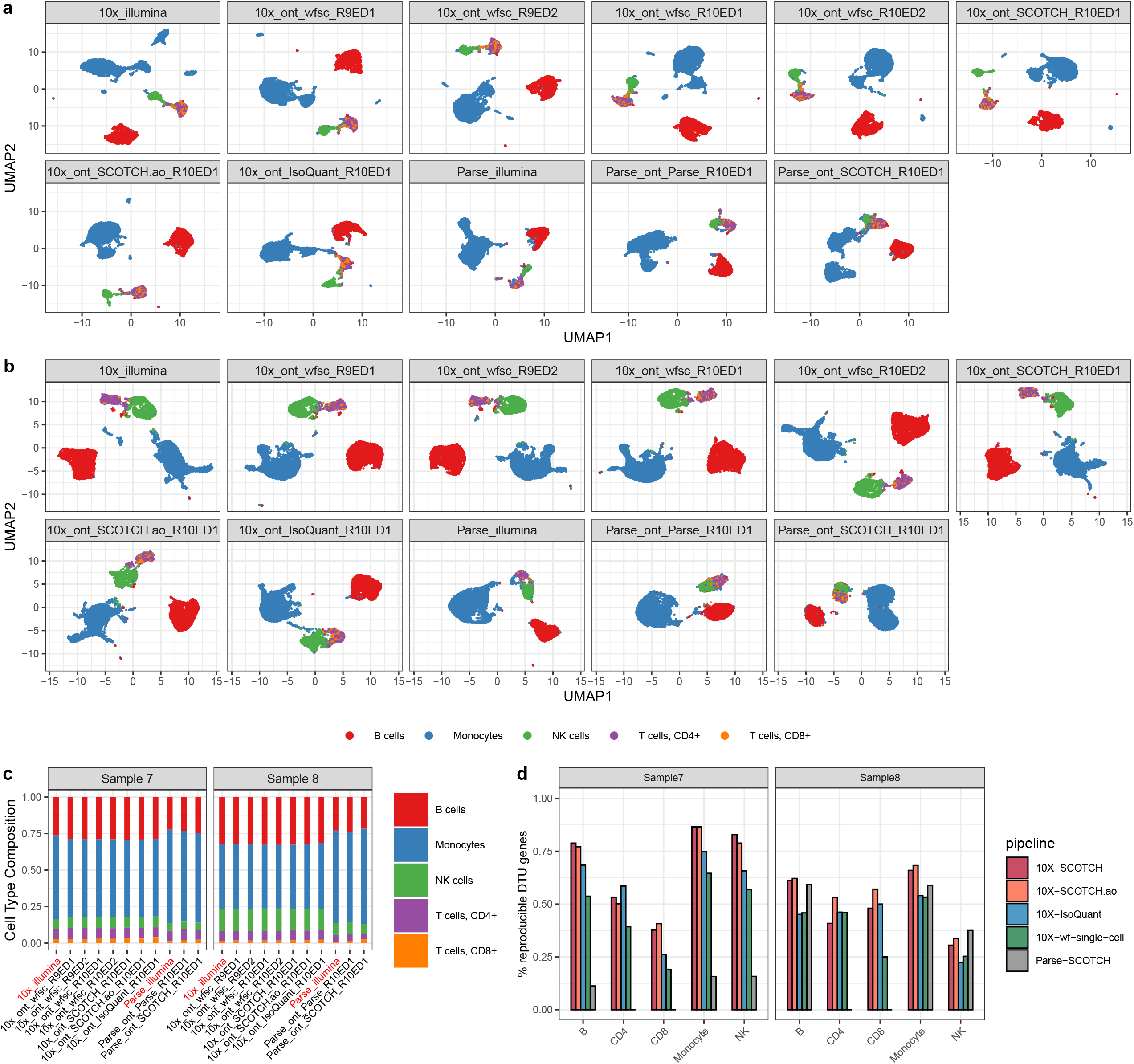
Consistency and reproducibility analysis comparing various technical platforms and pipelines. (a-b) UMAP visualization for clustering of different cell types comparing between 10X Genomics and Parse single-cell libraries, short-read and long-read scRNA-seq, R9 and R10 flowcells, Edit Distance of 1 and 2, as well as vendor (wf-single-cell, wfsc), SCOTCH pipelines, and IsoQuant on the same PBMC samples. Panel names are in the format of single-cell libraries_sequencing technology_computational pipeline_flowcell and edit distance. (a) PBMC sample 7. (b) PBMC sample 8. (c) Cell type compositions for sample 7 in a and sample 8 in b. (d) Percentage of DTU genes that are also identified in the other sample analyzed by SCOTCH and wf-single-cell pipelines for each cell type. DTU genes were identified using two-sided likelihood ratio tests with Holm-adjusted P values < 0.01 considered significant. Source data are provided as a Source Data file.

### SCOTCH aids in revealing transcriptome insights beyond gene expression

After confirming SCOTCH’s consistency and reproducibility across different platforms, we evaluated its ability to uncover transcript-level changes which conventional gene-level analysis may miss. To do this, we performed differential gene expression (DGE) analysis and differential transcript usage (DTU) analysis by comparing each cell type against the others using the gene-level, and transcript-level count matrices, respectively. We found that 1.5% to 26.7% of the genes, which were not differentially expressed between cell populations, exhibited DTU in both samples, while 6.3% to 51.6% exhibited DTU in at least one sample (**Figure 5a**). These findings suggest that relying solely on gene expression comparisons could overlook significant biological signals introduced by transcript variations. **Figure 5b** displays the number of DTU genes identified in both samples, highlighting differential transcript usage as a common feature among various cell types. Monocytes exhibited the highest number of DTU genes (n = 3731), with 1263 genes being unique to this cell type, suggesting that distinct isoforms could play roles specific to monocyte functions. B cells had the second-highest DTU count (n = 2441), with 1745 genes overlapping with monocytes, indicating shared transcriptomic characteristics and pathways (**Figure S4**). NK cells, along with CD4+, and CD8+ T cells had a lower number of DTU genes, likely due to their lower prevalence in the samples. These findings underscore the cell-type-specific nature of transcript usage. To further explore these patterns, we conducted gene set enrichment analysis to identify biological processes related to cell type-specific transcript usage (**Figure S4**). We observed that certain biological processes are consistently enriched in most cell types, such as ribosome and antigen processing and presentation. Monocytes and B cells show the most similar pathway enrichment patterns, reflecting the substantial overlap of DTU genes between them.

**Figure 5.**
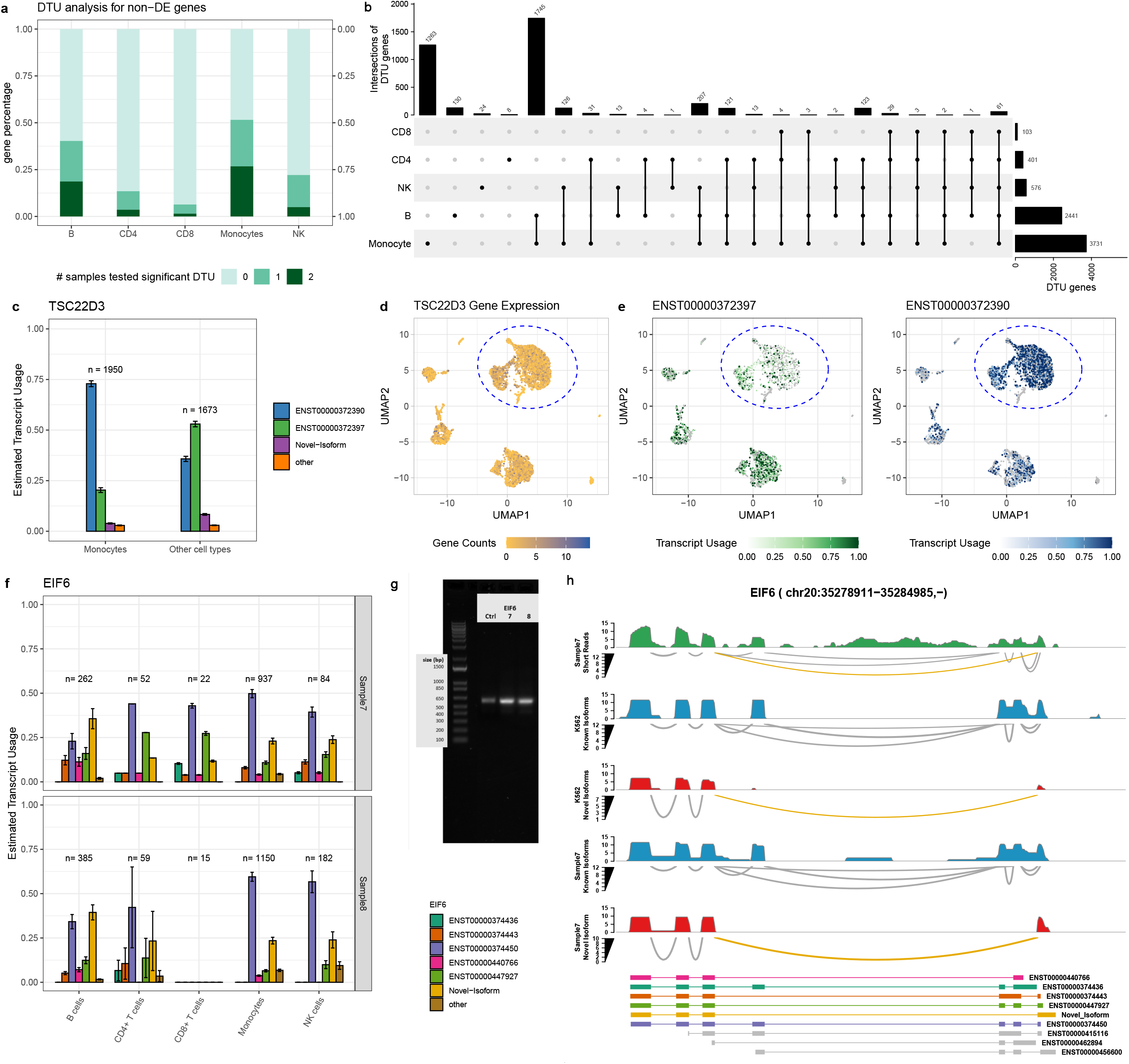
Transcript usage analysis for two PBMC samples using SCOTCH. (a) Proportions of non-differentially expressed (DE) genes with differential transcript usage (DTU) in various cell types. The darkness of colors indicates the number of samples in which significant DTU was observed. (b) Upset plot displaying the intersections of significant DTU genes of each cell type in against others in both sample 7 and sample 8. (c) Bar plot displaying the estimated average transcript usage for *TSC22D3 gene isofor*ms comparing monocytes and other cell types in sample 7. (d) UMAP visualization for expression levels of the *TSC22D3* gene in sample 7. (e) UMAP visualization for usages of ENST00000372397 and ENST00000372390 from the *TSC22D3* gene in each individual cell of sample 7. Cells within blue circles are monocytes. (f) Bar plot displaying estimated average transcript usages for *EIF6* gene isoforms comparing each cell type and other cell types for sample 7 and sample 8. (g) PCR validation of the *EIF6* novel isoform identified by SCOTCH. Novel isoform expression was compared in two independent PBMC samples (Sample 7 and Sample 8) and the K562 cell line (control, ctrl), with each lane representing one independent biological sample. Primers target the novel junction. (h) Displayed is sashimi plot of the *EIF6 gene* (chr20: 35,278,911–35,284,985, −) in sample7 and the K562 cell line, highlighting the presence of a novel isoform (orange) supported by short read. Coverage tracks are shown in green for short reads, blue for known isoforms in long reads, and red for novel isoforms in long reads. Splice junctions are represented in gray for known junctions and orange for novel junctions. In c and f, bars indicate mean estimated transcript usage across cells, and error bars represent model-derived standard deviations. n denotes cell numbers contributing to each group estimate. DTU genes were identified using two-sided likelihood ratio tests with Holm-adjusted P values < 0.01 considered significant. DE genes were identified using two-sided Wilconxon rank-sum tests with Holm-adjusted P values < 0.05 considered significant. Source data are provided as a Source Data file.

To showcase the utility of SCOTCH in identifying DTU genes, especially when overall gene expression levels remain unchanged, we examined the *TSC22D3* gene, which encodes the glucocorticoid-induced leucine zipper (GILZ) protein, a key regulator of anti-inflammatory and immunosuppressive responses^48–50^. GILZ modulates immune signaling by interfering with pathways such as NF-κB, AP-1, and Raf-MEK-ERK^51–53^, thereby influencing immune function and cell proliferation. As depicted in **Figure 5c-e**, transcript usage (TU) of *TSC22D3* significantly differs at both the gene level (p-adj < 0.0001) and the transcript level for ENST00000372390 (p-adj < 0.0001), ENST00000372397 (p-adj < 0.0001), ENST00000372383 (p-adj < 0.0001) and the novel-isoform (p-adj < 0.0001) between monocytes and other cell types in sample 7. Particularly, monocytes predominantly express ENST00000372390 (TU: 0.729±0.014) followed by ENST00000372397 (TU: 0.203±0.012), while in other cell types, ENST00000372397 becomes the predominant isoform (TU: 0.530±0.014), followed by ENST00000372390 (TU: 0.358±0.013). Despite these variations in transcript usage, overall *TSC22D3* expression levels remain unchanged between monocytes and other cell types, with a p-adj value of 0.922 (**Figure 5d**). These results suggest SCOTCH’s ability to uncover transcriptomic variations, even when gene expression levels show no significant changes.

We further examined the *EIF6* gene as another case study to demonstrate the utility of SCOTCH in DTU analysis. *EIF6* encodes a translation initiation factor that regulates ribosome assembly and plays critical roles in cell proliferation and tumorigenesis^54^. **Figure 5f** illustrates the DTU patterns for *EIF6* across five cell types in sample 7 and sample 8, comparing each cell type to others. SCOTCH identified a novel isoform expressed in all five cell types in both samples, except for CD8+ T cells in sample 8, likely due to the low abundance of this cell type. Among the known isoforms, ENST00000374450 dominates expression in four cell types, whereas the novel isoform is the predominant transcript in B cells. Notably, the novel isoform contains a novel splicing junction absent in previously annotated isoforms (**Figure 5h, Figure S7**). To validate its existence in two PBMC samples, we designed primers targeting the novel splicing junction and performed polymerase chain reaction (PCR). As shown in **Figure 5g**, PCR results confirmed the presence of this novel isoform not only in two PBMC samples but also in the K562 cell line, albeit at a lower abundance. We then validated the PCR band by Sanger sequencing, further confirmed the novel isoform sequence. We also quantified the fraction of reads mapped to the novel isoform: SCOTCH assigned 20.3% and 21.6% of total *EIF6* reads to the novel isoform in PBMC sample 7 and sample 8, respectively, while only 4.6% of reads were mapped to the novel isoform in the K562 cell line (**Figure 5h, Figure S7**). To further validate the novel junction, we analyzed both short-read and long-read data. We calculated the local relative junction abundance (LRJA), defined as the ratio of reads supporting the novel junction to reads supporting any junction involving the upstream exon. As shown in **Figure 5h** and **Figure S8**, the novel junction was confirmed in both short-read and long-read data is confirmed, with LRJA values of 0.111 and 0.190 for short-read sample 7 and sample 8, respectively, and 0.262 and 0.315 for their long-read counterparts. While the lower LRJA values in short-read data likely reflect its inherent limitation in capturing distant splice junctions, the consistent detection of the novel junction across both platforms provides strong validation for its existence. Together with the PCR results, these findings provide robust evidence for the novel isoform, further demonstrating the reliability of SCOTCH in identifying and annotating novel isoforms.

To further validate performance of SCOTCH on transcript-level quantification, we performed qPCR on FACS-sorted immune cell types, from two additional human PBMC samples. Primers were designed to target regions distinguishing ENST00000376049 and ENST000000376059 of the *AIF1* gene (**Figure 6a-b**). qPCR results confirmed dominant expression of ENST000000376059 in monocytes and NK cells, and of ENST00000376049 in CD4+ and CD8+ T cells (**Figure 6d**), consistent with SCOTCH quantifications for most cell types except B cells, which showed sample-specific differences (**Figure 6c**), and further supported by the long-read coverage profiles (**Figure 6a-b**). In addition to assessing transcript usage differences, SCOTCH elucidates isoform co-expression patterns using the mean-invariant over-dispersion parameter ϕ. Here we examined isoform usage patterns of the *AIF1* gene, which is instrumental in cytoskeletal rearrangements and cell migration^55^, and acts as a critical modulator of inflammation and immune cell activation^56^. To account for the imbalance in the numbers of cells expression *AIF1* isoforms, we down-sampled monocyte data to match the number of cells from other types, enabling a visually comparable analysis of cell type-specific isoform co-expression. **Figure S6** shows that monocytes typically express multiple isoforms simultaneously with similar usage proportions (Sample7: ϕ = 0.012, Sample8: ϕ = 0.010). In contrast, other cells exhibit greater heterogeneity, often exclusively expressing a single isoform, as indicated by higher dispersion values (Sample7 ϕ = 0.803, Sample8: ϕ = 0.529). These diverse isoform usage patterns have implications for the intricate roles of *AIF1* in cellular and immune processes.

**Figure 6.**
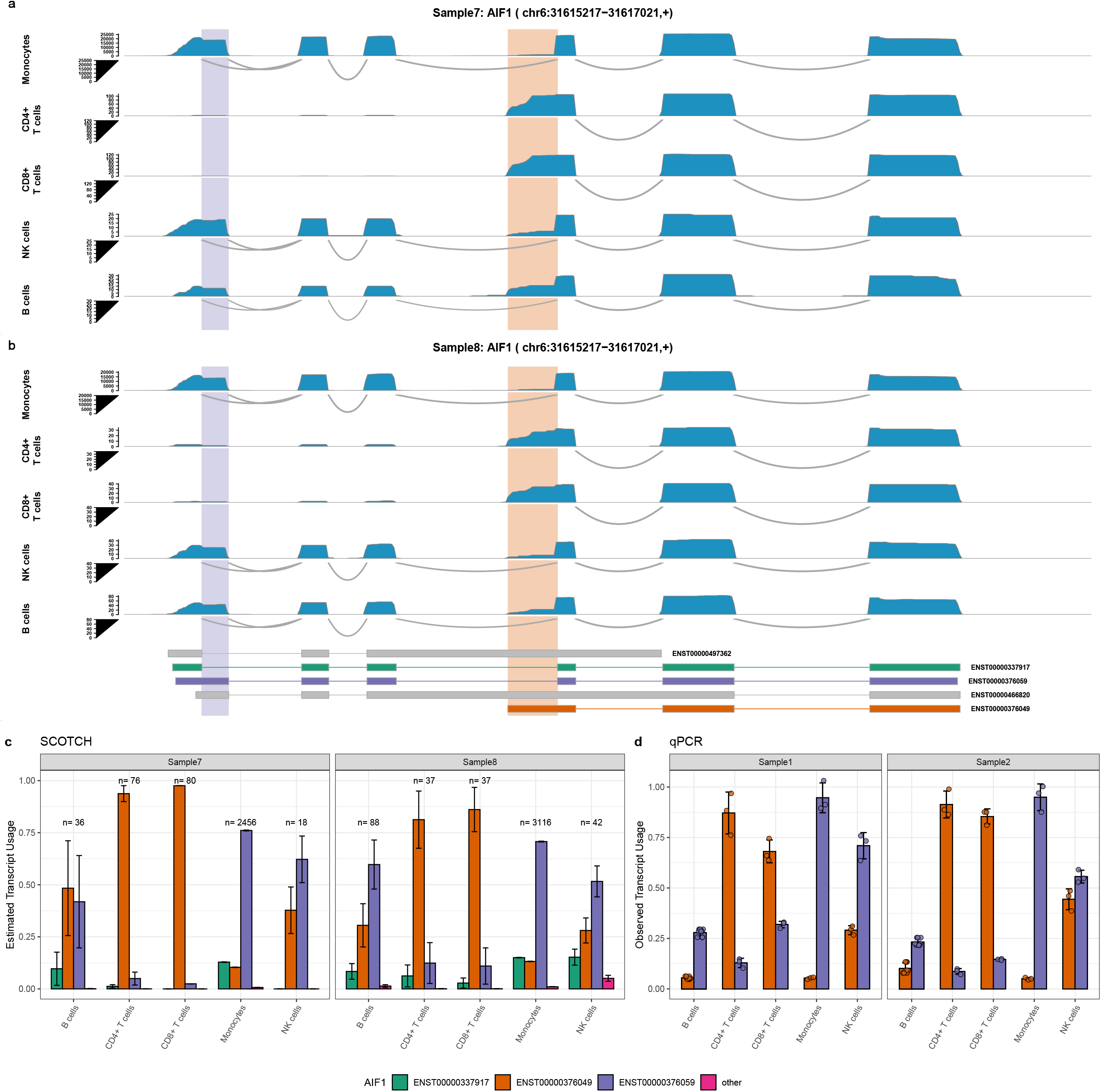
Cell type specific isoform usage of the *AIF1* gene (chr6: 31,615,217–31,617,021 +) across immune cell populations revealed by SCOTCH and validated by qPCR experiment. (a-b) Sashimi plots showing isoform expression of *AIF1* in immune cell types from Sample 7 and Sample 8 visualized by SCOTCH. The bottom panel displays known and novel isoforms of *AIF1*, with exon structures color-coded by transcript ID. Two highlighted regions (purple and orange) mark the positions targeted by primers used in qPCR validation to distinguish major isoforms of ENST00000376059 and ENST00000376049. (c) Estimated average transcript usage by SCOTCH across five immune cell types. Bars indicate mean estimated transcript usage across cells, and error bars represent model-derived standard deviations. n denotes cell numbers contributing to each group estimate. (d) Experimental validation of transcript usage by qPCR across sorted immune cell types in Sample 1 and Sample 2 for two major expressed isoforms. Points denote technical replicates from a single biological sample (n = 3 per group). Bars represent mean observed transcript usage, and error bars represent standard deviation. Source data are provided as a Source Data file.

### SCOTCH uncovers cell-type-specific transcript usage in human cerebral organoids

To investigate SCOTCH’s capability to uncover cell-type-specific transcript usage on the PacBio sequencing platform, we applied SCOTCH and IsoQuant to seven human cerebral organoid samples sequenced with the 10X-PacBio protocol^32^. Interestingly, we observed that the gene-level count matrix generated by IsoQuant is much sparser than that produced by SCOTCH (**Table S3**). Consequently, after preprocessing, SCOTCH retained a larger number of cells (n=4664) compared to IsoQuant (n=1406), enabling more accurate cell type annotations. As shown in **Figure 7a**, both methods produced similar proportions of progenitors and immature neurons. SCOTCH achieved a balanced proportion of neurons and progenitors, consistent with the findings from the original study. In contrast, IsoQuant’s sparse count matrix led to weak gene signature enrichment, making some clusters difficult to annotate. As a result, these clusters were labeled as other cell types due to insufficient gene signature specificity. When comparing transcript usage between neurons and progenitors, SCOTCH identified 194 DTU genes, while IsoQuant detected only 13 (**Figure 7b**), highlighting SCOTCH’s superior sensitivity in capturing cell-type-specific transcriptomic differences. The 194 DTU genes identified by SCOTCH are enriched in pathways related to cytoplasmic translation and substantia nigra development (**Figure 7d**). These genes are also enriched for postsynaptic density (PSD) proteins^57^ and targets of FMRP^58^ (**Figure S5**). Among the DTU genes, SCOTCH and IsoQuant both identified *CLTA* and *CLTB* as top 10 candidates, displaying consistent transcript usage patterns between the methods (**Figure 7c**). In progenitors, *CLTA* is predominantly expressed through the isoform ENST00000345519, whereas neurons exhibit a balanced usage of isoforms ENST00000396603 (neurons vs progenitors: SCOTCH: 0.323+0.089 vs 0.031+0.002, adjusted p-value < 0.0001; IsoQuant: 0.338 + 0.063 vs 0.030 + 0.000, adjusted p-value < 0.0001) and ENST00000345519 (neurons vs progenitors: SCOTCH: 0.350+0.093 vs 0.853+0.009, adjusted p-value < 0.0001; IsoQuant: 0.510+0.071 vs 0.933+0.000, adjusted p-value < 0.0001). For *CLTB*, neurons primarily express the isoform ENST00000310418 (neurons vs progenitors: SCOTCH: 0.913+0.000 vs 0.220+0.113, adjusted p-value < 0.0001; IsoQuant: 0.870+0.000 vs 0.243+0.080, adjusted p-value < 0.0001), while progenitors shift to isoform ENST00000345807 (neurons vs progenitors: SCOTCH: 0.087 +0.000 vs 0.780 +0.113, adjusted p-value < 0.0001; IsoQuant: 0.130+0.000 vs 0.757+0.080, adjusted p-value < 0.0001). The isoform usage patterns can also be observed in the read coverage plots (**Figure 7e-f**), where neurons and progenitors display distinct coverage profiles across specific exons for *CLTA* and *CLTB*. These DTU patterns are consistent with previous studies, where neuron-specific splicing of *CLTA* and *CLTB* influences synaptic vesicle cycling, neural differentiation, and brain development^11, 21, 30, 59, 60^.

**Figure 7.**
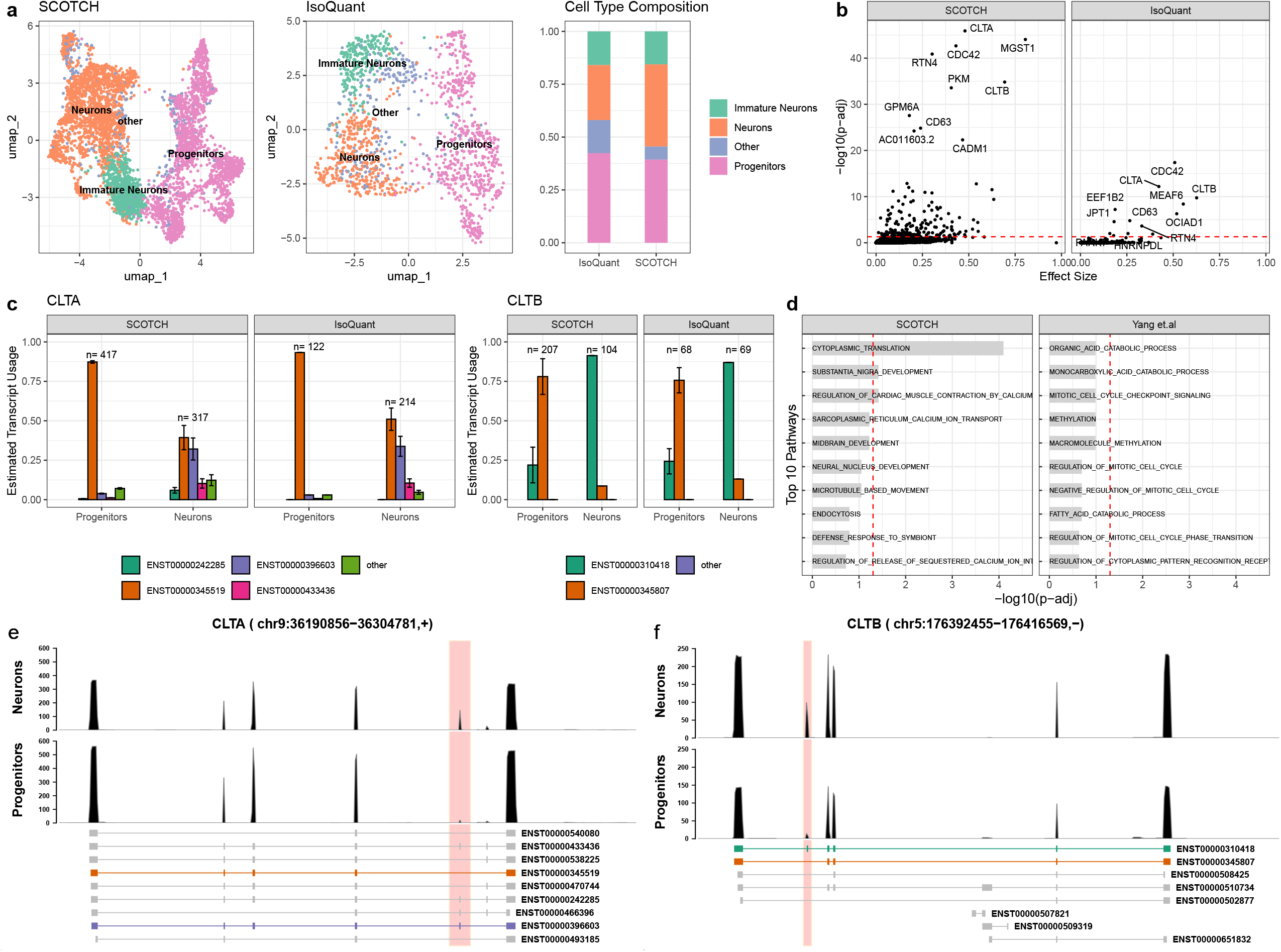
Transcript usage analysis on human cerebral organoid samples. (a). Displayed are UMAP visualization for cell type clusters generated by SCOTCH (left) and IsoQuant (middle), and corresponding cell type proportions (right). (b) Scatter plot showing DTU genes between neurons and progenitors detected by SCOTCH (left) and IsoQuant (right). The x-axis represents the effect size, defined as the maximum transcript usage difference between two cell types for each gene. Top 10 significant DTU genes are labeled. (c) Bar plots showing the estimated average transcript usage for genes *CLTA (left) and CLTB* (right) in neuron cells and progenitor cells. Bars indicate mean estimated transcript usage across cells, and error bars represent model-derived standard deviations. n denotes cell numbers contributing to each group estimate. (d) Top 10 enriched pathways for DTU genes identified by SCOTCH (left) and differential spliced genes identified by Yang et al^32^ (right). The x-axis represents negative log10-transformed adjusted P-values. (e) Displayed are reads mapped to *CLTA genes in ne*urons and progenitors. (f) Displayed are reads mapped to *CLTB genes in ne*urons and progenitors. In b and d, DTU genes were identified using two-sided likelihood ratio tests with Holm-adjusted P values < 0.05 considered significant. Pathway enrichment was evaluated using a one-sided hypergeometric test with Benjamini-Hochberg-adjusted P values < 0.05 considered significant. Red dashed lines indicate the significance threshold of adjusted P-value. In e and f, the most frequently used transcripts are colored in the annotation panel. The red-highlighted region marks exons with different splicing patterns between two isoforms. Error bars show standard deviations. Source data are provided as a Source Data file.

## Discussion

In this study, we generated a benchmarking dataset of two human PBMC samples and demonstrated the improved read quality of R10 flowcells over R9 flowcells through comparing five different sequencing approaches: 10X + Illumina, 10X + ONT_R9, 10X + ONT_R10, Parse + Illumina, and Parse + ONT_R10, under ED settings of 1 and 2 (**Table 1**). These advancements with R10 flowcells underscore the technical potential of long-read single-cell RNA sequencing (lr-scRNA-seq), enabling a shift in computational tool development from barcode correction to better unraveling transcriptome complexity. In response, we introduced SCOTCH, a suite of computational and statistical pipelines specifically designed for the processing and analysis of lr-scRNA-seq data. SCOTCH’s preprocessing pipeline is compatible with both Nanopore and PacBio sequencing platforms and supports single-cell libraries from 10X Genomics and Parse Biosciences. To the best of our knowledge, SCOTCH is the first publicly available tool that supports data from the Parse Biosciences platform, which offers a unique advantage by enabling simultaneous sequencing of multiple samples, eliminating the need for additional data integration and facilitating the analysis of specialized cell populations.

Despite recent advances in lr-scRNA-seq technologies, several challenges remain. First, ambiguous read-isoform mapping can occur at both the gene level and isoform levels, as reads may align to multiple overlapping or chimeric genes, or several transcripts of the same gene due to sequencing and alignment noise. Most existing tools do not resolve gene-level ambiguity prior to transcript-level assignment, where static thresholds are commonly used to handle read-isoform alignment and may introduce multiple mappings or mis-assignment errors. Ambiguous reads are often discarded or downweighted, leading to read loss, increased sparsity at both gene and transcript levels, and inaccurate quantifications (**Figure 3a, b, f, g, Figure 7a**). Second, accurate novel isoform identification requires a balance between sensitivity and specificity, ensuring recovery of true novel isoforms while minimizing false positives and redundant transcripts introduced by noise, which can distort downstream analyses and obscure genuine signals. Existing splice-graph–based approaches are highly susceptible to alignment artifacts, often producing fragmented or redundant transcript models, or alternatively sacrificing sensitivity when the graph is overly simplified (**Figure 3d, e, j**). Third, computational tools for preprocessing and downstream analysis remain fragmented, as most existing long-read isoform quantification tools were originally developed for bulk sequencing data and require manual adaption for single-cell application generated by different platforms. Finally, capturing transcript-level cell-to-cell heterogeneity is challenging because transcript-level count matrices are much sparser than gene-level ones. This problem is further exacerbated when existing tools discard large fractions of reads during isoform assignment, making the data even sparser (**Figure 3a, Figure 7a, Table S3**). Reliable biological insights therefore depend on accurate isoform identification and quantification with reliable statistical methods to separate true biological variability from technical noise.

SCOTCH incorporates several innovative design elements to address these limitations, making it highly effective for analyzing lr-scRNA-seq data. First, SCOTCH resolves ambiguous read-gene and read-isoform mappings through the incorporation of read mapping scores and a dynamic thresholding strategy that adapt to local noise patterns and ensure accurate assignment with minimal read loss (**Figure 3a, Figure 7a, Table S3, Supplementary notes**). Second, to accurately identify and annotate novel isoforms, SCOTCH refines sub-exon boundaries using reference annotations and read coverage, constructs read– exon matching profiles, and infers transcript structures through iterative Louvain clustering with read remapping. Unlike conventional splice-graph–based approaches, this strategy avoids the generation of excessive false-positive or redundant novel transcript models caused by noisy graphs, as well as reduced sensitivity in recovering true novel isoforms resulting from overly aggressive graph simplification, enabling accurate and sensitive novel isoform detection while remaining robust to read truncation, sequencing noise, and poly(A) stretch–related artifacts (**Figure 3d, e, j, Figure 5f-h, Supplementary notes**). Third, SCOTCH implements a unified preprocessing workflow that handles read demultiplexing and cell barcode parsing and supports simultaneous preprocessing of multiple samples within a study. It produces a consensus annotation file tailored to the experimental design and provides a statistical framework for isoform-level quantification and downstream analyses. Finally, SCOTCH’s isoform-based analyses provide clearer biological interpretation and have shown superior performance over exon-based approaches in detecting isoform-level variations.^21^ Together, these features establish SCOTCH as a robust and adaptable tool for transcriptome analysis in lr-scRNA-seq.

We also acknowledge several limitations inherent to the SCOTCH pipeline and our study. First, SCOTCH currently performs Louvain clustering on reads that cannot be mapped to any known isoforms of a gene and uses read clusters to generate candidate novel isoform annotations. While effective, this approach could be further refined by weighting reads based on alignment quality—an approach inspired by methods like LIQA ^42^—to improve the accuracy of novel isoform annotations by accounting for variations in read quality. Second, SCOTCH’s statistical framework currently performs one-way comparisons, focusing on mean shifts in transcript usage composition vectors between two cell populations. Extending this to Dirichlet regression would allow SCOTCH to incorporate additional covariates, such as batch effects, in complex study designs. This enhancement would enable simultaneous testing of both mean shifts and variance changes in transcript usage, making SCOTCH more adaptable to nuanced experimental setups. Moreover, a set of five samples was pooled by the Parse single-cell library construction approach and subsequently sequenced using R10 flowcells, yet only two of them are used in the current study. Given the relatively low sequencing depth per sample, direct comparisons between the Parse results and those from the 10X Genomics library cannot be completed in the current study. However, we have demonstrated consistency in the detection of differentially used transcripts (DTUs) when there is an adequate number of cells and sufficient coverage.

In the rapidly advancing field of single-cell and single-nucleus sequencing, current protocols and computational methodologies present several challenges but also offer significant opportunities for improvement. One major challenge is the analysis of full-length mRNA isoforms, which is essential for accurately profiling cellular subtypes and identifying transcript variants, such as alternative splicing and alternative polyadenylation sites ^14^. Emerging tools like SCOTCH, alongside improvements in vendor-supplied computational pipelines, are paving the way for deeper insights into complex gene expression patterns. Additionally, high dropout rates significantly impede the accuracy of gene expression profiling ^61^, an issue that is particularly pronounced in long-read transcriptome analysis at the transcript level. A promising future direction involves the development of computational methods specifically tailored for long-read data to mitigate this issue, building on approaches that have shown effectiveness in short-read data^62–64^. Furthermore, non-polyA RNA, which includes various functionally important regulatory RNAs ^65^, is better captured by the Parse Biosciences protocol due to its use of a mix of polyA primers and random hexamers, unlike the 10X Genomics protocol that requires a polyA tail exclusively. Future experimental approaches could further refine these techniques to provide a more comprehensive analysis of the transcriptome, ensuring that even non-polyadenylated RNA molecules are accurately represented. Additionally, future improvements are poised to refine unsupervised clustering utilizing full-length transcript information, leading to enhanced resolution at the transcript level and deeper biological insights. Another exciting frontier is the assessment of RNA velocity through long-read sequencing data ^66^, which holds the potential to drastically alter our understanding of RNA dynamics and regulatory mechanisms. Lastly, advancements in sequencing technology that increase depth and reduce costs are expected to facilitate more precise allele-and isoform-resolution counting, thereby broadening the applicability of transcriptomic analyses across diverse cell populations. This progression promises a more detailed and nuanced understanding of transcriptomics in health and disease. Collectively, these efforts are critical for refining lr-scRNA-seq analyses and advancing comprehensive cellular profiling.

In summary, by capturing the repertoire of transcriptional isoforms across diverse cell types, SCOTCH provides a foundation for delving into the functional significance of these isoforms in various cell types and holds promise in unraveling the complexity of cellular processes.

## Methods

### Preparation of the PBMC samples

Blood samples from two de-identified individuals, one male and one female, were collected using tubes coated with EDTA. These samples were promptly processed to separate PBMCs through Ficoll density gradient centrifugation at the Center for Applied Genomics (CAG) at the Children’s Hospital of Philadelphia (CHOP). The Institutional Review Board at the CHOP approved this study.

### Single cell library preparation on the 10X Genomics platform

RNA samples were processed using 10X Genomics Next GEM Single Cell 3’ Kit (V3.1) following manufacturer recommended protocols. During 10x library preparation, an aliquot of about 10ng of full-length amplified barcoded cDNA was taken for Nanopore sequencing. The remaining amount was processed further using the standard 10X protocol for Illumina sequencing.

### Single cell library preparation on the Parse Biosciences platform

RNA samples were processed using Parse Bioscience Evercode WT Mini v2 Kit following the manufacturer recommended protocol. During the first barcoding step, two control samples were distributed across 3 wells, Sample 7 and Sample 8 were distributed across 4 wells each, and a PBMC control was added to the last well. Approximately 70ng of each sublibrary was taken for Nanopore sequencing while remaining amount was used for Illumina sequencing.

### Oxford Nanopore sequencing of single-cell libraries

The single cell cDNA was processed for Nanopore sequencing using the Oxford Nanopore Technology (ONT) Ligation Sequencing Kit V14 (ONT, SQK-LSK114), cDNA-PCR Sequencing Kit (ONT, SQK-PCS111), or a combination of the cDNA-PCR Sequencing Kit and the Rapid Sequencing Kit (ONT, SQK-RAD114), and two custom primers, 5’-/5Biosg/CAGCACTTGCCTGTCGCTCTATCTTCCTACACGACGCTCTTCCGATCT-3’ and 5’-CAGCTTTCTGTTGGTGCTGATATTGCAAGCAGTGGTATCAA CGCAGAG-3’. The primers target the Illumina Read 1 sequence and the 10X Genomics template switch oligo sequence. 10ng of the single cell cDNA was mixed with 10 uM of both custom primers and LongAmp Hot Start Taq 2X Master Mix (NEB, M0533S). Amplification in a thermal cycler was performed using the following condition: 94C for 3 minutes, four cycles of 94C for 30 seconds, 66C down to 58C for 40 seconds, 58C for 50 seconds, and 65C for 6 minutes, a final extension cycle of 65C for 10 minutes, and a hold of 4C. Clean-up was performed using AMPure XP beads (Beckman Coulter™ cat # A63881) with a 1.25x solution to beads ratio, and washed twice with freshly made 200 ul of 80% ethanol, and the amplified cDNA was eluted with 10 ul of nuclease-free water.

Full-length cDNA was isolated using M280 streptavidin, 10 ug/ul (Invitrogen, 11205D), which the biotinylated cDNA binds to. 4 ml of a 2X wash/bind buffer was prepared with 10 mM Tris-HCl pH 7.5, 2 M NaCl, and 1 mM EDTA. Half of the 2X wash/bind buffer was used to make a 1X wash/bind buffer to us as a wash buffer. A 5 ug/ul streptavidin bead was made by replacing the buffer from 5 ul of the streptavidin beads with 10 ul of the 2X wash/bind buffer. 10 ul of the biotinylated cDNA was added to the 5 ug/ul beads and incubated at room temperature for 20 minutes. The mixture was then washed thrice with 1 ml of the 1X wash/bind buffer. A final wash was performed using 200 ul of 10 mM Tris-HCl pH 7.5. The beads were then resuspended in 20 ul of nuclease-free water.

The isolated full-length cDNA was then amplified with PCR. The 20 ul of the amplicon-bead conjugate was added to a 30 ul mixture of 10 uM PCR primer, LongAmp Hot Start Taq 2X Master Mix (NEB, M0533S), and nuclease-free water. The PCR primer used was dependent on the flow cell and/or sample used. For the R9 we used cPRM from SQK-PCS111, for the R10 for Sample7 and Sample8 we used cPRM from SQK-PCS111. Amplification in a thermal cycler was performed using the following condition: 94C for 3 minutes, four cycles of 94C for 15 seconds, 56C for 15 seconds, and 65C for 6 minutes, a final extension of 65C for 10 minutes, and a hold of 4C. Clean-up was performed using AMPure XP beads (Beckman Coulter™ cat # A63881) with a 1.25x solution to beads ratio, and washed twice with freshly made 200 ul of 80% ethanol, and the amplified cDNA was eluted with 15 ul of nuclease-free water.

For the samples processed with a PRM primer, an additional end-prep step needs to be performed before the adapter can be ligated to the full-length transcripts. 200 fmols from the previous step was mixed with the Ultra II End-prep Reaction Buffer and Ultra II End-prep Enzyme Mix (NEB, E7546) and incubated at 20C for 5 minutes and 65C for 5 minutes. Clean-up was performed as previously, but with a 1.0X sample-to-beads ratio and eluted with 60 ul of nuclease-free water.

For the R9 and Sample7 and Sample8 R10 samples Rapid Adapter T was added to 35 fmol of cDNA and incubated at room temperature for 5 minutes. For the remaining R10 samples, the end-prepped cDNA was added to a mixture of Ligation Buffer (LNB), NEBNext Quick T4 DNA Ligase, and Ligation Adapter (LA). The mixture was then incubated for 10 minutes at room temperature. Clean-up was done with a 2.5x sample-to-beads ratio, washed twice with 250 ul of the Short Fragment Buffer (SFB), and eluded with 25 ul of Elution Buffer.

The Parse samples were processed for Nanopore sequencing using the Ligation Sequencing Kit V14. The libraries were end repaired by mixing the Parse samples with the Ultra II End-prep Reaction Buffer and Ultra II End-prep Enzyme Mix and incubated at 20C for 5 minutes and 65C for 5 minutes. Clean-up was performed with a 1.0X sample-to-beads ratio and eluted with 60 ul of nuclease-free water. To add the Nanopore adapter to the cDNA, the end-prepped cDNA was added to a mixture of Ligation Buffer (LNB), NEBNext Quick T4 DNA Ligase, and Ligation Adapter (LA). The mixture was then incubated for 10 minutes at room temperature. Clean-up was done with a 2.5x sample-to-beads ratio, washed twice with 250 ul of the Short Fragment Buffer (SFB), and eluded with 25 ul of Elution Buffer.

Samples are loaded into an R9 or R10 PromethION flowcell (FLO-PRO002 or FLO-PRO114M) using the standard Nanopore loading protocol that involves priming the flow cell twice with a flow cell flush and flow cell tether mixture and then adding the Nanopore library that has been mixed the sequencing buffer and loading beads. Libraries were sequenced on a P2-solo and 96 hours.

### Validation of isoform expression by quantitative PCR

To validate novel isoform expression, we performed polymerase chain reaction (PCR) on cDNA generated from two human PBMC samples and commercially purchased total RNA of K562 cell line (Thermo Fisher Scientific, AM7832) to detect the *EIF6* novel isoform. The forward primer used was 5’-CAGCACAAACAGAGCAGGTTT-3’ and the reverse primer was 5’-GCTTGTTTACTGGTTACTTGGAGA-3’. PCR was performed using Invitrogen’s Platinum II Taq Hot-Start DNA polymerase at their standard 50ul protocol. The PCR product of the novel isoform was visualized using gel electrophoresis and the expected amplicon size is 588 bp. For cell-type-specific transcript quantification, we isolated CD4+ T cells, CD8+ T cells, monocytes, and natural killer (NK) cells from two additional human PBMC samples from UPENN’s Human Immunology Core. Total RNA was isolated from these samples by CHOP’s Center for Applied Genomics. Isoform expression was measured using quantitative PCR (qPCR). To distinguish between *AIF1* isoforms ENST00000376049 and ENST00000376059, isoform-specific primers targeting unique exonic regions were designed. The primers used to detect ENST00000376049 were 5’-CGTTGTCTCCTCCACCTAGC-3’ (forward) and 5’-AAGGGATAAGCGCTGACTGG-3’ (reverse), while those for ENST00000376059 were 5’-TGTCTCCCACCTCTACCAG-3’ (forward) and 5’-TGGAGGGGCAGATCCTCATCA-3’ (reverse). We used the absolute quantification method to quantify the isoforms, and standard curves were generated using the isoform’s primers. A stock solution of each isoform amplicon was generated with PCR with each primer pair in separate reactions, a mixture of the single cell samples was used as the input DNA. The amplicons where quantified using a Qubit Fluorometer and the number of copies were calculated from the concentration and amplicon size. Each individual stock solution was combined into one single standard stock. A 10X serial dilution of 6 points was then used as the standard curve for each primer pair. qPCR was performed using Applied Biosystems’ PowerTrack SYR Green Master Mix at the standard 10ul protocol with the fast-cycling method.

### Nanopore data processing

Sequencing data was basecalled using Guppy v6.5.7 with super-accuracy (SUP) model (also basecalled by Dorado, yielding similar results as shown in **Figure S12** and **Table S4**). After basecalling, cell UMI barcodes were identified using single cell library vendor supplied tools, including Nanopore’s wf-single-cell pipeline v1.0.1 (https://github.com/epi2me-labs/wf-single-cell) and Parse Bioscience’s pipeline. Since Parse Bioscience’s pipeline does not support transcript-level count matrix. Gene and transcript count matrices are generated using wf-single-cell, SCOTCH, and IsoQuant. To show improved accuracy of R10 flowcells over R9 flowcells, the pipeline was run twice with the R9 and R10 data for sample 7 and sample 8. Once with a barcode edit distance of 1 and a second time with a barcode edit distance of 2. Barcode edit distance is the allowance of a specified number of errors a barcode can have to be classified as a valid cell barcode. To use the Nanopore data on the Parse pipeline, the Nanopore reads would have to be split into artificial paired-end reads. With read 2 containing the Parse barcode sequences and read 1 containing the rest of the reads.

### scRNA-seq pre-processing, cell clustering, and cell type assignment

For PBMC samples, count matrices generated by different computational methods were processed using the Seurat package (v5.0.1)^67, 68^ in R (v4.3.1). Cells with detected genes below 200 or over the 99^th^ percentile or that had mitochondrial gene percentage over 20% were filtered out. The count matrices were then normalized, and the top 2000 variable genes were identified. Top 15 principal components were used to build the neighborhood graph, which was then clustered using the Louvain algorithm with resolution of 0.1. Cell subtyping was performed utilizing SingleR (v2.12.0) and the celldex::DatabaseImmuneCellExpressionData() function^69^. For visual representation and dimensionality reduction, Uniform Manifold Approximation and Projection (UMAP)^70^ was applied.

For raw bam files of cerebral organoid samples, we followed the same preprocessing pipeline and parameters as described in^32^. Count matrices generated by SCOTCH and IsoQuant were processed using the Seurat package. Cells with detected genes below 300 or over 5000 or that had mitochondrial gene percentage over 5% were filtered out. The count matrices were then normalized, and the top 2000 variable genes were identified. Top 30 principal components were used to build the neighborhood graph, which was then clustered using the Louvain algorithm with the resolution that can generate the same number of 22 cell clusters as the original study. To assign cell type labels, we used the AUCell package (v1.22.0)^71^ to calculate cell type-specific enrichment scores for each cell cluster based on marker genes identified in the original study. Cells were then assigned to their respective cell types.

### The SCOTCH pipeline for isoform level characterization of single cell RNA-seq data

The SCOTCH pipeline requires two types of input files: a long-read RNA-seq file provided as a tagged BAM file, and an optional isoform annotation file in GTF format. Upon processing these inputs, SCOTCH generates count matrices at both the gene level and the transcript level, with the capability to identify novel isoforms do not present in the original annotation file. Leveraging these gene and transcript count matrices, SCOTCH enables statistical analyses of the transcriptome across different cell populations, including the detection of genes with differential isoform usage and the identification of isoform switching events. SCOTCH is available at https://github.com/WGLab/SCOTCH. Detailed description of the computational methods is given below.

1. Prepare annotated bam files: Depending on the library type, we utilize either the ‘wf-single-cell’ or ‘IsoSeq’ pipeline for 10X Genomics libraries or ‘Parse’ for Parse libraries to perform read alignment and barcode identifications. This process produces BAM files annotated with barcode information.
2. Prepare gene annotation files: SCOTCH supports three modes for this step: annotation-only, annotation-free and enhanced-annotation mode. In annotation-only mode, exons are segmented into non-overlapping sub-exons solely based on GTF gene annotation files. In enhanced-annotation mode, read coverage data from BAM files is integrated to refine sub-exon annotations to improve novel isoform identification. This process involves filtering low-coverage regions, identifying splicing positions, and detecting sharp changes in read coverage to discover new exons and more accurately partition existing exons into sub-exons. In annotation-free mode, gene and exon structures are inferred entirely from read coverage data (see the **Methods** section on *Exon discovery procedure in enhanced-annotation mode and annotation-free mode*). Detailed gene information, including gene name, strand direction, genome locations, sub-exon coordinates, and transcript annotations, is compiled and stored in a pickle file. Additionally, overlapping genes are grouped into non-overlapping meta-genes for more accurate read-to-gene and read-to-isoform mapping.
3. Assign reads to annotated isoforms: Gene isoforms are treated as specific different combinations of non-overlapping sub-exons. A read is considered aligned to a sub-exon if it covers more than 60% of the sub-exon’s length, and is deemed to have skipped the sub-exon if it fails to cover more than 20% of its length, otherwise as ambiguously alignment. Initially, reads are mapped to all annotated isoforms, and any read-isoform pairing is excluded if: (1) the read covers sub-exons do not present in the annotated isoform, or (2) the read skips sub-exons that are present in the annotated isoform. Notably, possible read truncations or degradations are not regarded as skipping sub-exons. We specifically implement a dynamic thresholding strategy to handle small-sized sub-exons during read-isoform mapping. Starting from a minimal threshold, we increase it in 10 bp increments until it reaches 80 bp or average sub-exon length of the gene, whichever is smaller. This process continues until we achieve the first unique mapping or, if none is found, the first multiple mapping. Small sub-exons are excluded from decisions on whether a read maps to an isoform to enhance resilience against erroneous mappings and reduce ambiguous alignments. If a read still aligns with multiple annotated isoforms, we select the isoform with the fewest unmapped exons. To further accommodate specific analysis needs, users have the flexibility to adjust parameters of sub-exon coverage and lengths thresholds.
4. Identify novel isoforms: Following previous steps, if a read cannot be mapped to any existing annotated isoforms, it is classified as a ‘novel read’ to be assigned to a novel isoform. To identify novel isoforms, we construct a read-read similarity graph with a maximum of 1500 novel reads at a time. Each node represents a read, and edge weights between nodes are determined by their shared sub-exon mapping patterns. Using the Louvain method, we detect read communities that likely represent the same novel isoforms. For each read community, we determine the novel isoform annotation by calculating the mode of sub-exon mapping encodings across all reads. Once a set of novel isoform candidates is generated, all remaining unmapped reads are realigned to these candidates. Any remaining unmapped reads are grouped into a new batch of 1500, and the process is repeated until no further graphs can be constructed. After generating all novel isoform candidates, novel reads are remapped to the identified isoforms using a relaxed threshold for small sub-exons (the smaller value between 80 bp or the average sub-exon length) to encourage multiple mappings. Redundant isoforms are then eliminated if reads mapped to them can also map to other isoforms, refining the final set of novel isoforms.
5. Handle read ambiguity: Except in cases where a read may map to multiple isoforms, for which we reduce ambiguity through dynamic thresholding, reads may map to several overlapping or non-overlapping genes (e.g. chimeric reads due to sequencing artifacts). To resolve these ambiguities, we adopt a mapping score strategy to determine both read-gene and read-isoform assignments. The mapping score is calculated as the percentage of bases aligned to an isoform. Reads are prioritized as follows: first, to genes where the read maps to an existing annotated isoform; second, to genes corresponding to a novel isoform; and lastly, to genes where the read aligns with exons but cannot be classified as either known or novel isoforms.
6. Generate count matrix: During count matrix construction, novel isoforms are retained if supported by reads exceeding a user-defined pseudo-bulk threshold, defined either as an absolute read count or relative transcript usage within the gene (default: absolute count ≥ 20). Isoforms below this threshold are classified as non-categorized novel. Cell barcodes are then used to generate gene- and transcript-level count matrices.
7. Transcriptome analysis: SCOTCH can perform transcriptome analysis by testing differential transcript usage on both gene and transcript levels and identify isoform switching events. For gene *g* with *K* isoforms, we model the transcript expression level *X*_*ck*_ for isoform *k* in cell *c* using a Dirichlet-multinomial hierarchical model to account for transcript usage variations among cells. We assume *X*_*ck*_ ~ MultiNomial (*n*_c_,π_*c*_), and π_*c*_ Dirichlet (α). Here, 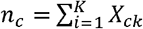 is the total transcript counts for the gene in the cell. π_*c*_ = (π_*c*,1_ … π_*c,k*_) is cell-specific transcript usage. α = (α_1_,…, α _K_) are the concentration parameters of the Dirichlet distribution we will estimate, and the average transcript usage for the cell population is 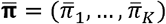 with 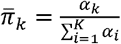. Similar with LongCell, we define 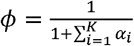 as a mean-invariant over-dispersion parameter, representing inter-cell heterogeneity. A small *ϕ* suggests that cells are likely to express various isoforms with similar usage proportions across the cell population, and a large *ϕ* value indicates a more exclusive expression manner, with each cell predominantly expressing one isoform while different cells may express others. Our goal is to estimate 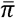 using maximum likelihood estimation and compare between different cell populations *A* and *B* using likelihood ratio test. On the gene level, we test 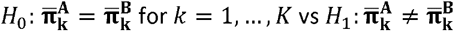.for any 1 ≤ *k* ≤ *K*. On the transcript level, we test 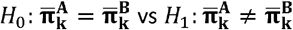 for isoform k. To mitigate the impact of low-abundance isoforms on statistical inference, rare isoforms may optionally be aggregated into one isoform using user-defined thresholds, and novel isoforms may also be combined into a unified novel category for downstream analysis. We define isoform switching events if the dominant isoform except the aggregated rare isoforms is different between two cell populations inferred by 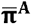 and 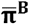. The effect size is defined as 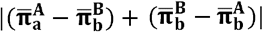 where isoform *a* and *b* are dominant isoforms for cell populations *A* and *B*, respectively.

### Exon discovery procedure in enhanced-annotation mode and annotation-free mode

In the enhanced-annotation mode of SCOTCH, we refine sub-exon annotations by incorporating read coverage data from BAM files. First, we extract read coverage across genome regions of interest based on existing gene annotation files. Regions with coverage below 2% of the maximum coverage (coverage_threshold_exon) or fewer than 20 reads are excluded to retain high-confidence coverage regions. We then identify contiguous coverage blocks, representing likely meta-exons, and remove blocks shorter than 20bp to eliminate unreliable segments. Next, we partition exon blocks into sub-exons utilizing both splicing junction and coverage change patterns as described following. 1. Splicing junctions: We analyze splicing positions by counting the number of reads supporting each splice junction. Positions with fewer than 2% of the maximum splicing count (coverage_threshold_splicing) or fewer than 20 reads are discarded. Splicing positions located within 10 bp of each other are merged to avoid over-segmentation, and those within 10 bp of known exon boundaries are filtered out to minimize noise. 2. Coverage change patterns: We smooth the read coverage data using a Gaussian filter and calculate the first derivative to identify local changes. Z-scores are computed from these derivative values, and positions where the absolute z-score exceeds 10 (z_score_threshold) are flagged as sharp changes, potentially indicating novel transcript start or end sites. To prevent over-partitioning, sharp changes that occur more than 10 bp away from known exon boundaries are excluded. By applying these steps, we are able to refine sub-exon annotations.

For annotation-free mode, novel gene regions are first identified based solely on coverage data, and then similar procedures are applied to partition these regions into sub-exon segments using splicing junctions and coverage change patterns.

### Simulation study

We simulated ground-truth nanopore scRNA-seq data for two distinct cell populations using a modified version of lrgasp-simulation (https://github.com/andrewprzh/lrgasp-simulation/tree/main). While originally developed for bulk long-read RNA-seq data, we adapted the simulation tool for single-cell applications by introducing between-cell transcript usage variations. We used GENCODE annotations for all 1080 genes that have at least two known isoforms on chromosome 6. To simulate the discovery of novel isoforms, we randomly removed 2466 (30%) isoforms and designated them as ground-truth novel isoforms. The remaining 5762 (70%) isoforms, covering all 1080 genes, were used as the existing annotation file for input across all tools. For a given gene *g*, let *K* represent the true number of isoforms based on the reference annotation file. We sampled gene expression counts *n*_*c*_ for the cell *c* from the distribution *n*_*c*_ ~ Poisson(*λ*_*c*_), where *λ*_*c*_ ~ *U* (0,15) We then sampled the true transcript usage for the cell from the distribution π_*c*_ ~ Dirichlet(α), where each α_*k*_ is drawn from a Gamma (2, 2) distribution. The expected transcript counts for cell ***c*** are *x*_*c*_ = (*x*_*c*,1_, *x*_*c*,2,_ … *x*_*c,k*_), where *x*_*c,k*_ = *n*_*c*_ π_*c,k*_ Based on transcript lengths, transcript per million (TPM) were calculated as input for the lrgasp-simulation. To control gene counts and underlying ground truth isoform usage, we simulated reads for each cell expressing one gene independently. We simulated 1000 cells for each of the two populations, with 540 genes sharing the same *α* values and the remaining 540 having different a values to simulate differential transcript usage (DTU) between populations. In total, 15,381,985 reads were simulated across the two populations. The simulated FASTA files were combined and aligned to the human genome using minimap2^72^. Cell barcodes and UMIs were added as tags in the BAM files based on the cell origins of the reads.

## Data availability

Raw data of two PBMC samples using both 10X Genomics and Parse libraries, Illumina and nanopore sequencings have been deposited to Sequence Read Archive (SRA) at the National Center for Biotechnology Information (NCBI) under accession number PRJNA1105904. The human cerebral organoids data used in this study can be downloaded from^32^. Source data are provided with this paper.

## Code availability

The SCOTCH software developed in this study is publicly available at https://github.com/WGLab/SCOTCH under the MIT license. Version 1.0.0 of SCOTCH has been archived in Zenodo and is accessible via DOI: http://doi.org/10.5281/zenodo.18841221.

## Acknowledgements

This study is supported by NIH grant GM132713, HG013359 and the CHOP Research Institute. We thank the IDDRC Biostatistics and Data Science core (NIH grant HD105354) for technical support on machine learning and high-performance computing.

## Author Contributions Statement

J.C. performed the sequencing experiments, participated in data analysis and processing. Z.X. designed the computational software and benchmarked its performance through biological samples. S.M. participated in simulation study data analysis. H.Q. performed clustering data analysis, visualization and interpretation, and participated in writing the manuscript. C.K. advised on experimental design, generated the 10X Genomics and Parse libraries and coordinated the study. H.H. provided insights into technologies, guided experimental design and guided data interpretation. K.W. conceived and supervised the study. All authors read and participated in writing the manuscript.

## Competing Interests Statement

The authors declare no competing interests.

## Notes

### Competing Interest Statement

The authors have declared no competing interest.

### Summary of Updates

The Figure 3-7 are updated, and the relevant description in Results are updated.

## References

1. Jovic, D. et al. Single-cell RNA sequencing technologies and applications: A brief overview. Clin Transl Med 12, e694 (2022).

2. Ke, M., Elshenawy, B., Sheldon, H., Arora, A. & Buffa, F.M. Single cell RNA-sequencing: A powerful yet still challenging technology to study cellular heterogeneity. Bioessays 44, e2200084 (2022).

3. Wang, S. et al. The Evolution of Single-Cell RNA Sequencing Technology and Application: Progress and Perspectives. Int J Mol Sci 24 (2023).

4. Conte, M.I., Fuentes-Trillo, A. & Dominguez Conde, C. Opportunities and tradeoffs in single-cell transcriptomic technologies. Trends Genet 40, 83–93 (2024).

5. Ake, F. et al. Quantification of transcript isoforms at the single-cell level using SCALPEL. Nat Commun 16, 6402 (2025).

6. Wang, Y., Zhao, Y., Bollas, A., Wang, Y. & Au, K.F. Nanopore sequencing technology, bioinformatics and applications. Nat Biotechnol 39, 1348–1365 (2021).

7. van Dijk, E.L. et al. Genomics in the long-read sequencing era. Trends Genet 39, 649–671 (2023).

8. Marx, V. Method of the year: long-read sequencing. Nat Methods 20, 6–11 (2023).

9. Warburton, P.E. & Sebra, R.P. Long-Read DNA Sequencing: Recent Advances and Remaining Challenges. Annu Rev Genomics Hum Genet 24, 109–132 (2023).

10. Park, E., Pan, Z., Zhang, Z., Lin, L. & Xing, Y. The Expanding Landscape of Alternative Splicing Variation in Human Populations. Am J Hum Genet 102, 11–26 (2018).

11. Vaquero-Garcia, J. et al. A new view of transcriptome complexity and regulation through the lens of local splicing variations. Elife 5, e11752 (2016).

12. Rogalska, M.E., Vivori, C. & Valcarcel, J. Regulation of pre-mRNA splicing: roles in physiology and disease, and therapeutic prospects. Nat Rev Genet 24, 251–269 (2023).

13. Marasco, L.E. & Kornblihtt, A.R. The physiology of alternative splicing. Nat Rev Mol Cell Biol 24, 242–254 (2023).

14. Gupta, I. et al. Single-cell isoform RNA sequencing characterizes isoforms in thousands of cerebellar cells. Nat Biotechnol (2018).

15. Singh, M. et al. High-throughput targeted long-read single cell sequencing reveals the clonal and transcriptional landscape of lymphocytes. Nature communications 10, 3120 (2019).

16. Rebboah, E. et al. Mapping and modeling the genomic basis of differential RNA isoform expression at single-cell resolution with LR-Split-seq. Genome biology 22, 1–28 (2021).

17. Hahaut, V. & Picelli, S. Full-Length Single-Cell RNA-Sequencing with FLASH-seq. Methods Mol Biol 2584, 123–164 (2023).

18. Shi, Z.X. et al. High-throughput and high-accuracy single-cell RNA isoform analysis using PacBio circular consensus sequencing. Nat Commun 14, 2631 (2023).

19. Al’Khafaji, A.M. et al. High-throughput RNA isoform sequencing using programmed cDNA concatenation. Nat Biotechnol 42, 582–586 (2024).

20. Lebrigand, K. et al. The spatial landscape of gene expression isoforms in tissue sections. Nucleic Acids Res 51, e47 (2023).

21. Joglekar, A. et al. A spatially resolved brain region- and cell type-specific isoform atlas of the postnatal mouse brain. Nat Commun 12, 463 (2021).

22. Li, H. et al. Improving gene isoform quantification with miniQuant. Nat Biotechnol (2025).

23. Fu, Y. et al. Single cell and spatial alternative splicing analysis with long read sequencing. BioRxiv (2023).

24. Tian, L. et al. Comprehensive characterization of single-cell full-length isoforms in human and mouse with long-read sequencing. Genome Biol 22, 310 (2021).

25. Lebrigand, K., Magnone, V., Barbry, P. & Waldmann, R. High throughput error corrected Nanopore single cell transcriptome sequencing. Nat Commun 11, 4025 (2020).

26. Wang, Q. et al. Single cell transcriptome sequencing on the Nanopore platform with ScNapBar. RNA 27, 763–770 (2021).

27. Chen, C. et al. Single-cell multiomics reveals increased plasticity, resistant populations, and stem-cell-like blasts in KMT2A-rearranged leukemia. Blood 139, 2198–2211 (2022).

28. Sereika, M. et al. Oxford Nanopore R10.4 long-read sequencing enables the generation of near-finished bacterial genomes from pure cultures and metagenomes without short-read or reference polishing. Nat Methods 19, 823–826 (2022).

29. Ni, Y., Liu, X., Simeneh, Z.M., Yang, M. & Li, R. Benchmarking of Nanopore R10.4 and R9.4.1 flow cells in single-cell whole-genome amplification and whole-genome shotgun sequencing. Comput Struct Biotechnol J 21, 2352–2364 (2023).

30. Joglekar, A. et al. Single-cell long-read sequencing-based mapping reveals specialized splicing patterns in developing and adult mouse and human brain. Nat Neurosci 27, 1051–1063 (2024).

31. Byrne, A. et al. Single-cell long-read targeted sequencing reveals transcriptional variation in ovarian cancer. Nat Commun 15, 6916 (2024).

32. Yang, Y. et al. Single-cell long-read sequencing in human cerebral organoids uncovers cell-type-specific and autism-associated exons. Cell Rep 42, 113335 (2023).

33. Penter, L. et al. Integrative genotyping of cancer and immune phenotypes by long-read sequencing. Nat Commun 15, 32 (2024).

34. Shiau, C.K. et al. High throughput single cell long-read sequencing analyses of same-cell genotypes and phenotypes in human tumors. Nat Commun 14, 4124 (2023).

35. Chen, Y. et al. Context-aware transcript quantification from long-read RNA-seq data with Bambu. Nat Methods 20, 1187–1195 (2023).

36. Prjibelski, A.D. et al. Accurate isoform discovery with IsoQuant using long reads. Nat Biotechnol 41, 915–918 (2023).

37. Kabza, M. et al. Accurate long-read transcript discovery and quantification at single-cell, pseudo-bulk and bulk resolution with Isosceles. Nat Commun 15, 7316 (2024).

38. Blondel, V.D., Guillaume, J.-L., Lambiotte, R. & Lefebvre, E. Fast unfolding of communities in large networks. Journal of Statistical Mechanics: Theory and Experiment 2008 (2008).

39. Gandal, M.J. et al. Transcriptome-wide isoform-level dysregulation in ASD, schizophrenia, and bipolar disorder. Science 362 (2018).

40. Tekath, T. & Dugas, M. Differential transcript usage analysis of bulk and single-cell RNA-seq data with DTUrtle. Bioinformatics 37, 3781–3787 (2021).

41. Marques-Coelho, D. et al. Differential transcript usage unravels gene expression alterations in Alzheimer’s disease human brains. NPJ Aging Mech Dis 7, 2 (2021).

42. Hu, Y. et al. LIQA: long-read isoform quantification and analysis. Genome Biol 22, 182 (2021).

43. You, Y. et al. Identification of cell barcodes from long-read single-cell RNA-seq with BLAZE. Genome Biol 24, 66 (2023).

44. Tytgat, O. et al. Nanopore Sequencing of a Forensic STR Multiplex Reveals Loci Suitable for Single-Contributor STR Profiling. Genes 11 (2020).

45. González-Recio, O. et al. Sequencing of SARS-CoV-2 genome using different nanopore chemistries. Applied microbiology and biotechnology 105, 3225–3234 (2021).

46. Frankish, A. et al. GENCODE: reference annotation for the human and mouse genomes in 2023. Nucleic Acids Res 51, D942–D949 (2023).

47. Pertea, G. & Pertea, M. GFF Utilities: GffRead and GffCompare. F1000Res 9 (2020).

48. Cannarile, L. et al. Cloning, chromosomal assignment and tissue distribution of human GILZ, a glucocorticoid hormone-induced gene. Cell Death Differ 8, 201–203 (2001).

49. Li, Y. et al. TSC22D3 as an immune-related prognostic biomarker for acute myeloid leukemia. iScience 26, 107451 (2023).

50. D’Adamio, F. et al. A new dexamethasone-induced gene of the leucine zipper family protects T lymphocytes from TCR/CD3-activated cell death. Immunity 7, 803–812 (1997).

51. Ayroldi, E. et al. Modulation of T-cell activation by the glucocorticoid-induced leucine zipper factor via inhibition of nuclear factor kappaB. Blood 98, 743–753 (2001).

52. Mittelstadt, P.R. & Ashwell, J.D. Inhibition of AP-1 by the glucocorticoid-inducible protein GILZ. J Biol Chem 276, 29603–29610 (2001).

53. Ayroldi, E. et al. Glucocorticoid-induced leucine zipper inhibits the Raf-extracellular signal-regulated kinase pathway by binding to Raf-1. Mol Cell Biol 22, 7929–7941 (2002).

54. Miluzio, A. et al. Impairment of cytoplasmic eIF6 activity restricts lymphomagenesis and tumor progression without affecting normal growth. Cancer Cell 19, 765–775 (2011).

55. Autieri, M.V., Kelemen, S.E. & Wendt, K.W. AIF-1 is an actin-polymerizing and Rac1-activating protein that promotes vascular smooth muscle cell migration. Circulation research 92, 1107–1114 (2003).

56. Liu, G., Ma, H., Jiang, L. & Zhao, Y. Allograft inflammatory factor-1 and its immune regulation. Autoimmunity 40, 95–102 (2007).

57. Bayes, A. et al. Characterization of the proteome, diseases and evolution of the human postsynaptic density. Nat Neurosci 14, 19–21 (2011).

58. Darnell, J.C. et al. FMRP stalls ribosomal translocation on mRNAs linked to synaptic function and autism. Cell 146, 247–261 (2011).

59. Louadi, Z. et al. Functional enrichment of alternative splicing events with NEASE reveals insights into tissue identity and diseases. Genome Biol 22, 327 (2021).

60. Blue, R.E., Curry, E.G., Engels, N.M., Lee, E.Y. & Giudice, J. How alternative splicing affects membrane-trafficking dynamics. J Cell Sci 131 (2018).

61. Badsha, M.B. et al. Imputation of single-cell gene expression with an autoencoder neural network. Quant Biol 8, 78–94 (2020).

62. Van Dijk, D. et al. Recovering gene interactions from single-cell data using data diffusion. Cell 174, 716–729. e727 (2018).

63. Huang, M. et al. SAVER: gene expression recovery for single-cell RNA sequencing. Nature methods 15, 539–542 (2018).

64. Li, W.V. & Li, J.J. An accurate and robust imputation method scImpute for single-cell RNA-seq data. Nature communications 9, 997 (2018).

65. Picelli, S. et al. Full-length RNA-seq from single cells using Smart-seq2. Nat Protoc 9, 171–181 (2014).

66. Bergen, V., Soldatov, R.A., Kharchenko, P.V. & Theis, F.J. RNA velocity-current challenges and future perspectives. Mol Syst Biol 17, e10282 (2021).

67. Butler, A., Hoffman, P., Smibert, P., Papalexi, E. & Satija, R. Integrating single-cell transcriptomic data across different conditions, technologies, and species. Nature biotechnology 36, 411–420 (2018).

68. Satija, R., Farrell, J.A., Gennert, D., Schier, A.F. & Regev, A. Spatial reconstruction of single-cell gene expression data. Nature Biotechnology 33, 495–502 (2015).

69. Aran, D. et al. Reference-based analysis of lung single-cell sequencing reveals a transitional profibrotic macrophage. Nature immunology 20, 163–172 (2019).

70. Becht, E. et al. Dimensionality reduction for visualizing single-cell data using UMAP. Nature biotechnology 37, 38–44 (2019).

71. Aibar, S. et al. SCENIC: single-cell regulatory network inference and clustering. Nat Methods 14, 1083–1086 (2017).

72. Li, H. Minimap2: pairwise alignment for nucleotide sequences. Bioinformatics 34, 3094–3100 (2018).

