## Supplementary Materials for "SCOTCH: isoform-level characterization of gene expression through long-read single-cell RNA sequencing"

#### Supplementary tables and figures

**Table S1. Summary statistics of the single-cell PBMC datasets across sequencing and processing platforms. For some measures on Nanopore R9 flowcells, the statistic for using edit distance of 1 or 2 (ED1/ED2) are listed. For Illumina data and Nanopore R10 data, only ED1 was used.**

| Single Cell Platform | Sequencing Platform (Sample ID) | Sequence Saturation (ED1/ED2) | UMI Count Threshold (ED1/ED2) | Mean Reads Per Cell (ED1/ED2) | Median Genes Per Cell (ED1/ED2) | Median Transcripts Per Cell (ED1/ED2) | Median UMI Count Per Cell (ED1/ED2) |
|---|---|---|---|---|---|---|---|
| 10X | Illumina (Sample 7) | 71.7% | 500 | 51,542 | 2,723 | --- | 8,376 |
| 10X | ONT-R9 (Sample 7) | 34.7% / 35.8% | 507 / 565 | 11,983 / 13,570 | 2,279 / 2,398 | 1,280 / 1,360 | 6,954 / 7,724 |
| 10X | ONT-R10 (Sample 7) | 46.0% / 46.6% | 597 / 625 | 16,350 / 17,375 | 2,520 / 2,581 | 1,426 / 1,467 | 7,761 / 8,203 |
| 10X | Illumina (Sample 8) | 65.7% | 500 | 37,145 | 2,597 | --- | 6,893 |
| 10X | ONT-R9 (Sample 8) | 28.6% / 29.5% | 531 / 579 | 8,760 / 9,866 | 2,060 / 2,177 | 1,097 / 1,173 | 5,493 / 6,096 |
| 10X | ONT-R10 (Sample 8) | 43.9% / 44.4% | 587 / 631 | 14,398 / 15,219 | 2,471 / 2,528 | 1,359 / 1,397 | 6,967 / 7,320 |
| Parse | Illumina (Sample 7) | 62.3% | --- | 70,992 | 2,338 | 5,078 | --- |
| Parse | ONT-R10 (Sample 7) | 20.7% | --- | 7,129 | 1,116 | 1,758 | --- |
| Parse | Illumina (Sample 8) | 62.0% | --- | 32,903 | 1,407 | 2,282 | --- |
| Parse | ONT-R10 (Sample 8) | 20.0% | --- | 3,144 | 563 | 742 | --- |

**Table S2. Read assignment of different isoform categories by SCOTCH for Nanopore R10 data.**

| | Sample7<br>n (%) | Sample8<br>n (%) |
|---|---|---|
| **Known Isoforms** | 15943660 (55.45) | 18581863 (51.53) |
| **Novel Isoforms** | 11419114 (39.72) | 15387988 (42.67) |
| **Uncategorized** | 5914052 (20.57) | 2092857 (5.8) |
| **Total Reads** | 28752280 | 36062708 |

**Table S3. Total number of transcripts in the human cerebral organoid PacBio dataset processed by SCOTCH and IsoQuant**

| Sample | Total Reads | |
|---|---|---|
| | SCOTCH | IsoQuant |
| 84d_1 | 1464039 | 929218 |
| 84d_2 | 1891211 | 1226664 |
| 137d_1 | 2153486 | 1514764 |
| 137d_2 | 2329708 | 1651855 |
| 145d_1 | 1286609 | 894020 |
| 145d_2 | 699358 | 474287 |
| 145d_3 | 1447376 | 1006666 |

**Table S4. Summary statistics comparing basecaller of Guppy and Dorado**

| | | # Reads | # Bases | # Reads PF | # Bases PF | Q-score | N50 |
|---|---|---|---|---|---|---|---|
| **Sample7 R10** | **Guppy** | 199,157,241 | 161,314,352,872 | 168,592,908 | 133,906,287,373 | 18.96 | 943 |
| | **Dorado** | 199,133,412 | 155,870,701,157 | 138,389,424 | 119,566,540,623 | 11.19 | 942 |
| **Sample7 R9** | **Guppy** | 214,103,012 | 172,187,282,026 | 140,381,804 | 116,333,029,267 | 11.86 | 960 |
| | **Dorado** | 213,632,426 | 170,242,429,611 | 123,276,129 | 107,608,120,683 | 10.23 | 953 |
| **Sample8 R10** | **Guppy** | 231,670,493 | 176,538,129,203 | 204,670,670 | 152,668,392,018 | 19.66 | 876 |
| | **Dorado** | 231,641,804 | 171,132,110,366 | 172,081,920 | 137,787,173,003 | 11.45 | 871 |
| **Sample8 R9** | **Guppy** | 225,365,300 | 166,925,340,829 | 151,510,326 | 114,206,090,495 | 12.02 | 878 |
| | **Dorado** | 224,884,474 | 165,187,882,530 | 132,392,128 | 104,927,197,755 | 10.33 | 870 |

PF = Mean Q-score >=10

**Figure S1.** Influence of edit distance (ED) criteria on cell number, gene number, and transcript number for 10X Nanopore PBMC samples using R9 and R10 flowcells.

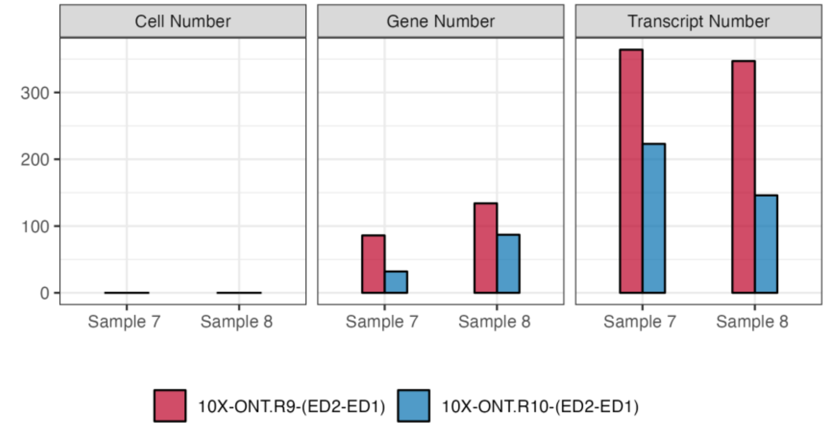


**Figure S2.** Venn diagrams showing gene numbers with significant differential transcript usage (DTU) that are consistent between sample7 and sample8 identified by different computational tools in each of the five cell types, compared against all other cells. DTU genes were identified using two-sided likelihood ratio tests with Holm-adjusted P values < 0.01 considered significant.

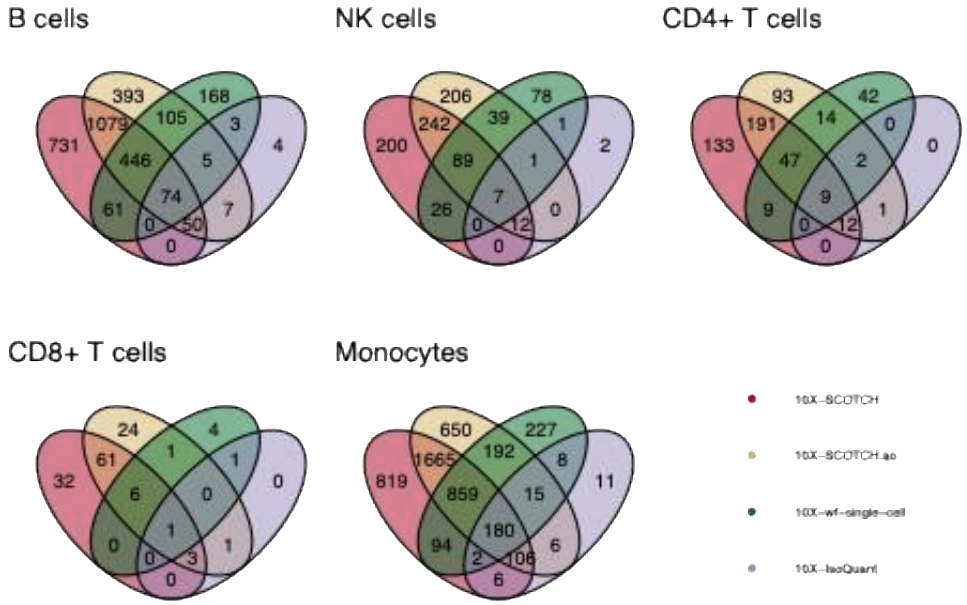

**Figure S3.** Venn diagrams showing the number of genes with significant differential transcript usage (DTU) that are consistent between sample 7 and sample 8 identified by SCOTCH for platforms of Parse Bioscience and 10X Genomics in each of the three cell types, compared against all other cells. DTU genes were identified using two-sided likelihood ratio tests with Holm-adjusted P values < 0.01 considered significant.

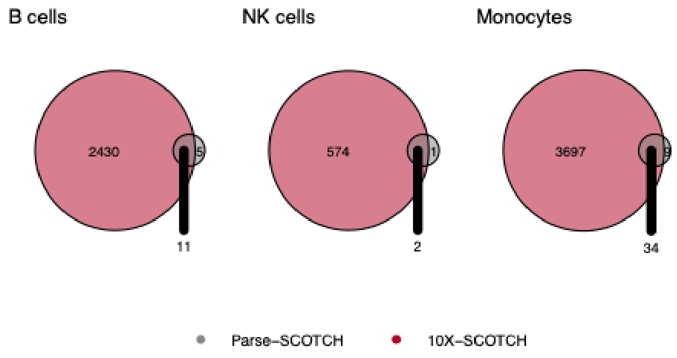


**Figure S4.** Pathway enrichment analysis of significant DTU genes for PBMC samples comparing each of the five cell types against all other cell types. x-axis indicates normalized enrichment score (NES). The darkness of colors indicates the significance level of enrichment analysis. DTU genes were identified using two-sided likelihood ratio tests with Holm-adjusted P values < 0.01 considered significant. Benjamini-Hochberg-adjusted P values < 0.05 was considered significant in pathway enrichment analysis.

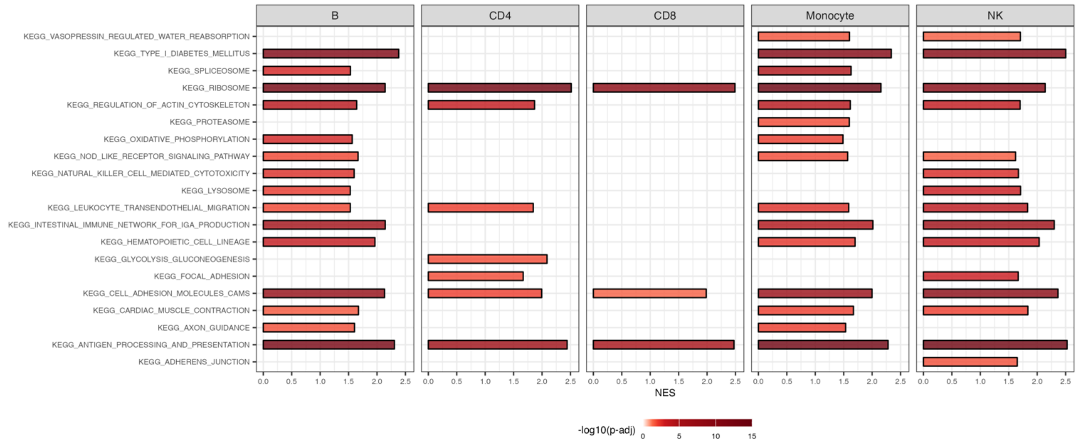

**Figure S5.** Comparison of enrichment analysis between DTU and DSE gene sets in human cerebral organoids dataset. SCOTCH-identified differential transcript usage (DTU) genes and Yang et al.'s differential spliced (DSE) genes were tested for enrichment in FMRP and hPSD gene sets. Odds ratios, confidence intervals, and p-values are shown. DTU genes were identified using two-sided likelihood ratio tests with Holm-adjusted P values < 0.05 considered significant.

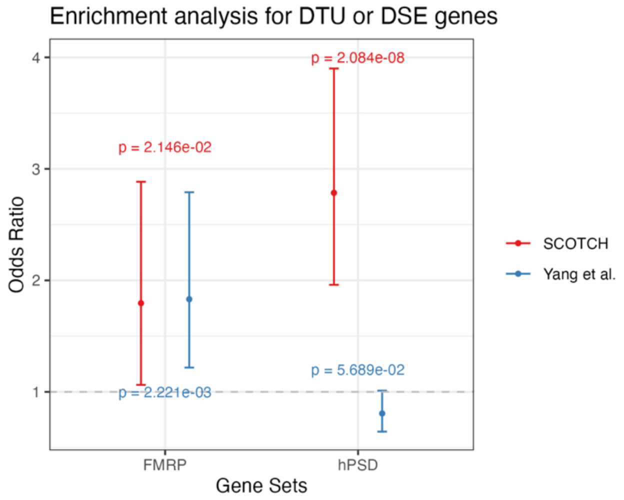


**Figure S6.** Transcript usage distribution of *AIF1* gene comparing monocytes and all other cell types for sample 7 and sample 8 of PBMC dataset. Each column represents a cell, with colors indicating isoform compositions used within each cell. Monocytes are downsampled to the same number of other cell types.

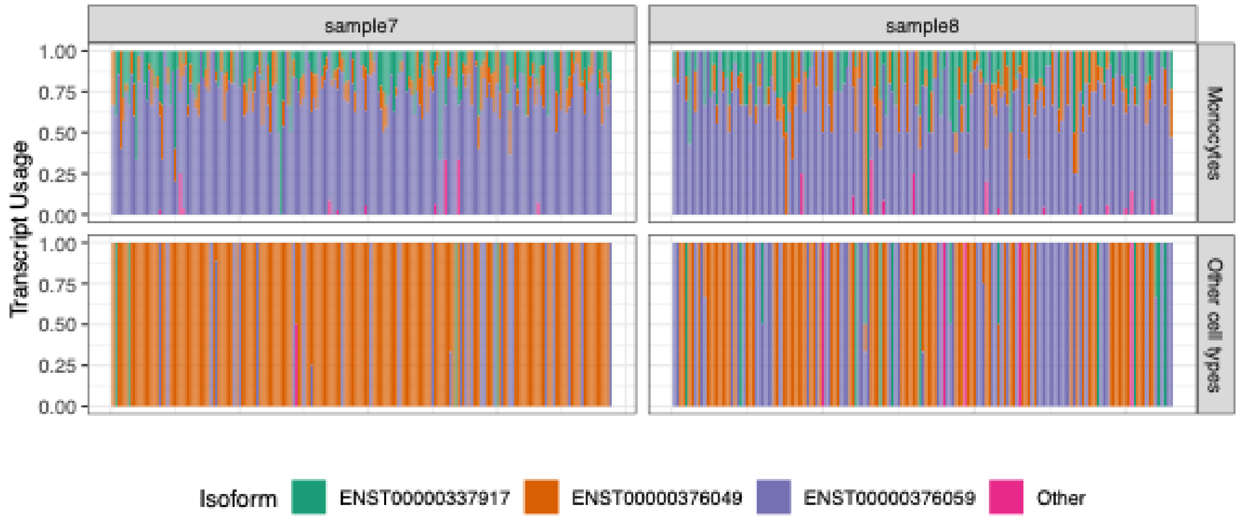

**Figure S7.** Displayed is sashimi plot of the *EIF6* gene (chr20: 35,278,911–35,284,985, - ) in sample8 of PBMC dataset and the K562 cell line, highlighting the presence of a novel isoform (orange) supported by short read. Coverage tracks are shown in green for short reads, blue for known isoforms in long reads, and red for novel isoforms in long reads. Splice junctions are represented in gray for known junctions and orange for novel junctions.

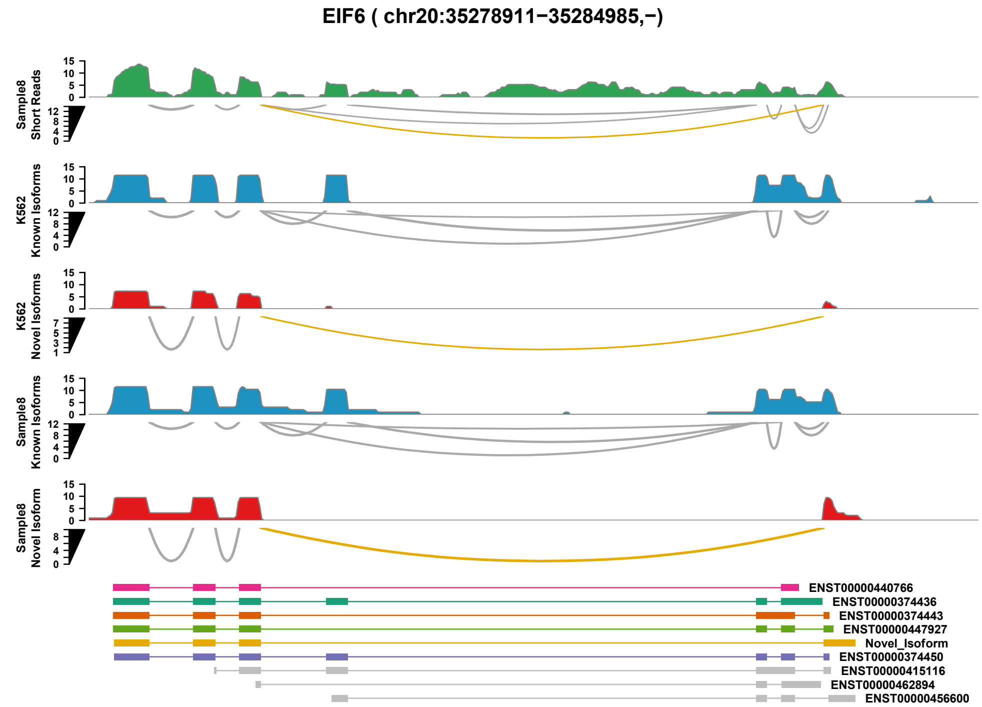


**Figure S8.** Validation of the novel splice junction using short-read and long-read data. Stacked bar plots show the local relative junction abundance (LRJA) involving the upstream (3') exon of the novel splice junction in sample 7 and sample 8, based on long-read (lr) and short-read (sr) sequencing.

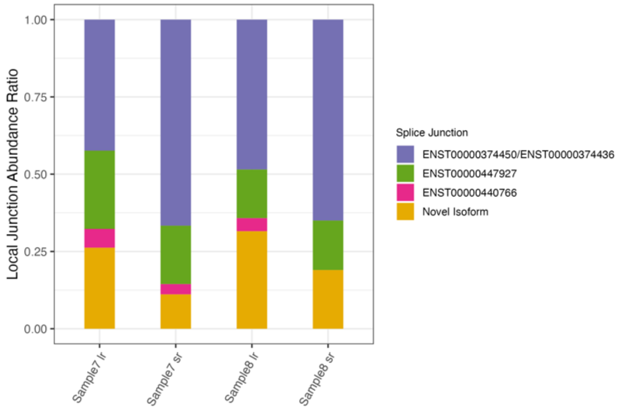

**Figure S9.** Knee plots for filtering barcodes across sequencing platforms for Sample 7. UMI counts (y-axis) are plotted against ranked barcodes (x-axis) for sequencing platforms of Illumina and Oxford Nanopore Technologies (ONT). Valid barcodes (blue) are distinguished from background (gray).

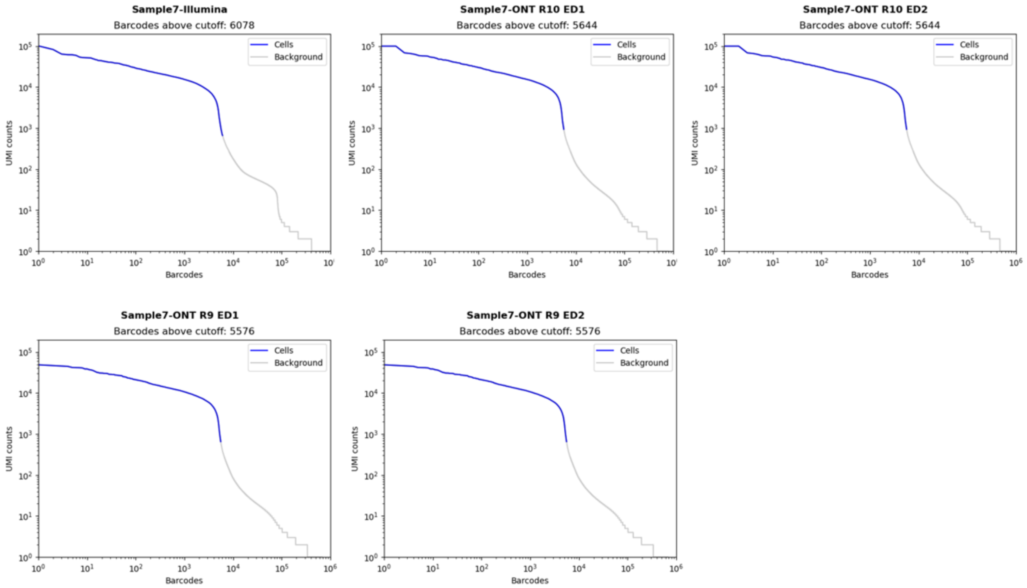


**Figure S10.** Knee plots for filtering barcodes across sequencing platforms for Sample 8. UMI counts (y-axis) are plotted against ranked barcodes (x-axis) for sequencing platforms of Illumina and Oxford Nanopore Technologies (ONT). Valid barcodes (blue) are distinguished from background (gray).

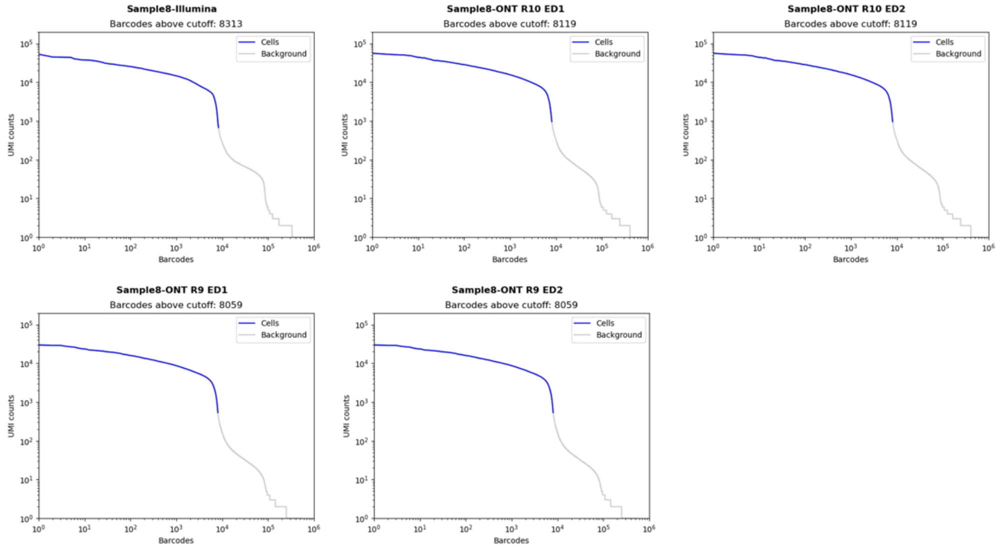

**Figure S11.** Knee plots for filtering barcodes across sequencing platforms for Sample 7 and Sample 8. UMI counts (y-axis) are plotted against ranked barcodes (x-axis) for sequencing platforms of Illumina and Parse Biosciences. Valid barcodes (blue) are distinguished from background (gray).

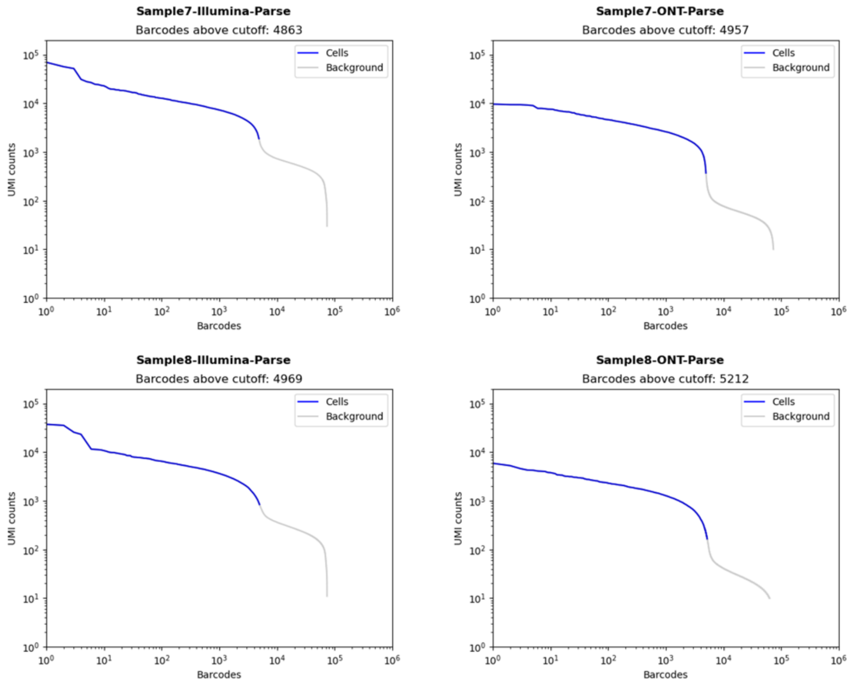

**Figure S12.** Influence of edit distance (ED) criteria on final results of five single-cell library + sequencing platforms generated by vendor computational pipeline: 10X + Illumina, Parse + Illumina, 10X + Nanopore_R9, 10X + Nanopore_R10, Parse + Nanopore R10. Cell number, total gene number, total transcript number, median number of genes per cell, and median number of UMI per cell that are identified based on different flowcell versions and different ED thresholds (1 or 2). Dorado basecaller was used.

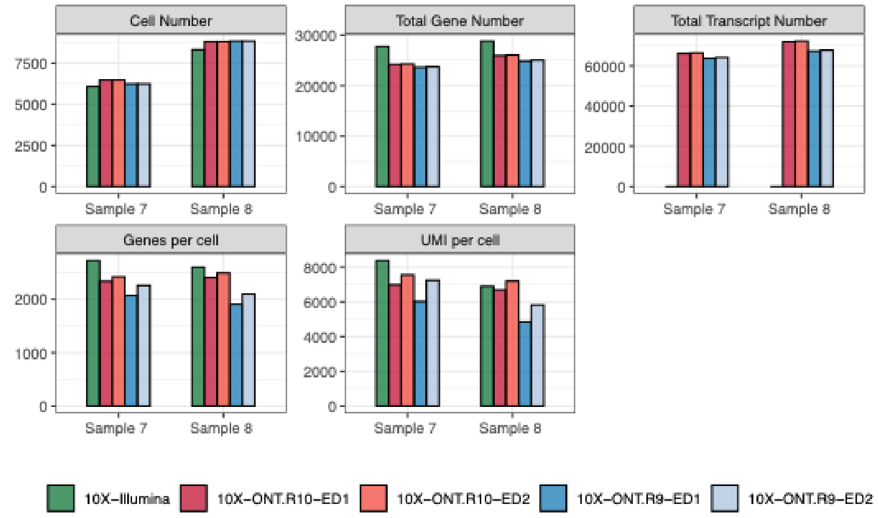

**Figure S13.** Comparison between exon-centric and splice-graph-based strategies in read-isoform mapping. **(a)** Enhanced-annotation mode of SCOTCH to refine existing exon boundaries using read coverage patterns. SCOTCH utilize curve patterns of overlaid exons (red dashed line and blue dashed line shaped a sharp coverage change indicated by green line) to decompose the rightmost exon of a known isoform (blue) into three non-overlapping sub-exon candidates (10, 11, and 12), enabling novel isoform identification with novel exon boundaries. Right panels illustrate exon-centric read–exon mapping using two alignment thresholds (*match_low* and *match_high*) and dynamic small-exon filtering (default 0–80 bp) to stabilize read–isoform compatibility. **(b)** Schematic representation of two annotated isoforms on the positive strand. In splice-graph representation, splice junctions (S1–S5) and transcript terminal sites (T1–T3) are treated as nodes, with exons serving as edges connecting nodes. In the exon-centric representation, isoforms are decomposed into non-overlapping sub-exons (E1–E7), and each isoform can be expressed as a combination of these sub-exons. **(c–g)** Representative scenarios illustrating differences between splice-graph–based and exon-centric mapping under increasing alignment complexity. **(c)** Read of isoform 2 with no minimal noise. **(d)** Read of isoform 2 with two consecutive exon boundary drifts. **(e)** Read of isoform 2 with a short unannotated exon combined with elongated exon. **(f)** Read of isoform 2 with short unannotated exon near the transcript start site co-occurs with read truncation. **(g)** Read of isoform 1 with junction shift, exon skipping, and terminal perturbation co-occur. Red-labeled segments (seg1–seg3) denote alignment artifacts. Blue and green profiles denote correct assignments to corresponding isoform. Black profiles indicate incorrect mappings, with red elements marking the source of artifact-induced errors

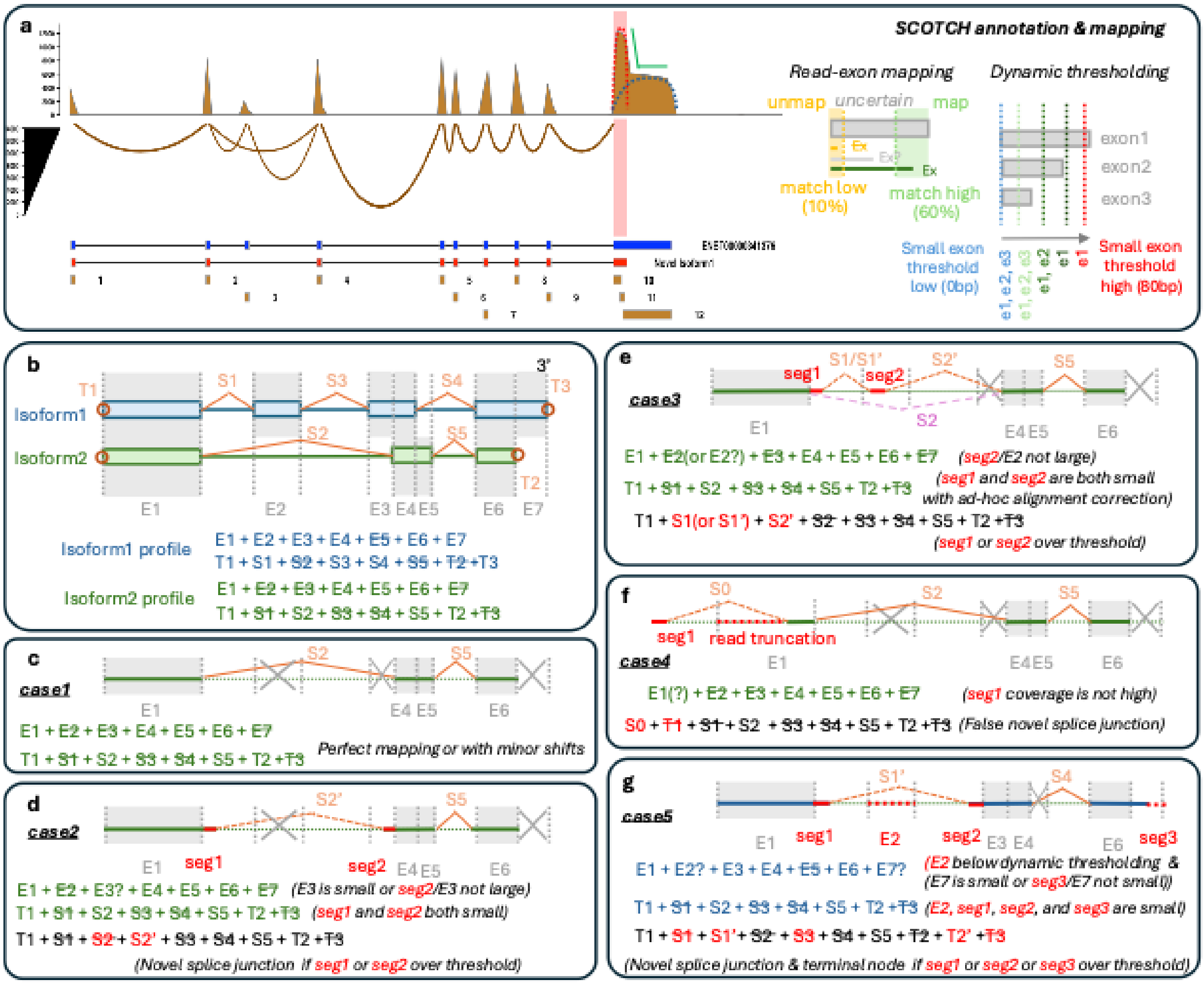

SCOTCH annotation & mapping
Read-exon mapping
Dynamic thresholding
unmap
uncertain
map
match low (10%)
match high (60%)
exon1
exon2
exon3
Small exon threshold low (0bp)
Small exon threshold high (80bp)
Isoform1
Isoform2
Isoform1 profile
E1 + E2 + E3 + E4 + E5 + E6 + E7
T1 + S1 + S2 + S3 + S4 + S5 + T2 +T3
Isoform2 profile
E1 + E2 + E3 + E4 + E5 + E6 + E7
T1 + S1 + S2 + S3 + S4 + S5 + T2 +T3
case1
E1 + E2 + E3 + E4 + E5 + E6 + E7
T1 + S1 + S2 + S3 + S4 + S5 + T2 +T3
Perfect mapping or with minor shifts
case2
E1 + E2 + E3? + E4 + E5 + E6 + E7 (E3 is small or seg2/E3 not large)
T1 + S1 + S2 + S3 + S4 + S5 + T2 +T3 (seg1 and seg2 both small)
T1 + S1 + S2 + S2' + S3 + S4 + S5 + T2 +T3
(Novel splice junction if seg1 or seg2 over threshold)
case3
E1 + E2(or E2?) + E3 + E4 + E5 + E6 + E7 (seg2/E2 not large)
T1 + S1 + S2 + S3 + S4 + S5 + T2 +T3 (seg1 and seg2 are both small with ad-hoc alignment correction)
T1 + S1(or S1') + S2' + S2 + S3 + S4 + S5 + T2 +T3
(seg1 or seg2 over threshold)
case4
read truncation
E1(?) + E2 + E3 + E4 + E5 + E6 + E7 (seg1 coverage is not high)
S0 + T1 + S1 + S2 + S3 + S4 + S5 + T2 +T3 (False novel splice junction)
case5
E1 + E2? + E3 + E4 + E5 + E6 + E7? (E2 below dynamic thresholding & (E7 is small or seg3/E7 not small))
T1 + S1 + S2 + S3 + S4 + S5 + T2 +T3 (E2, seg1, seg2, and seg3 are small)
T1 + S1 + S1' + S2 + S3 + S4 + S5 + T2 + T2' +T3
(Novel splice junction & terminal node if seg1 or seg2 or seg3 over threshold)

### Supplementary Notes

##### Robust read-isoform mapping in sub-exon space through dynamic thresholding

Long-read RNA sequencing data contain various types of artifacts arising from sequencing and alignment processes[1], including extra exons, exon skipping, exon boundary drift, poly-A/T stretch interference, and read truncations. In complex genes, these artifacts frequently co-occur within the same read. An effective isoform identification method must therefore distinguish true novel isoform signals from technical noise, rather than amplify such artifacts into spurious transcript models. The standard isoform reconstruction workflow typically proceeds in two stages: (1) mapping reads to existing annotated isoforms, and (2) constructing novel isoform annotations from unmapped alignments, followed by repeating the previous read-isoform mapping step. Importantly, read–isoform mapping is central to both stages of this process. Errors or instability in mapping decisions propagate throughout the workflow, directly affecting both the identification of known isoforms and the accuracy of novel isoform reconstruction.

Most existing methods rely primarily on splice-graph–based strategies. Even when exon information is incorporated (e.g., IsoQuant integrates splice and exon signals), the mapping decision is largely driven by splice junction detection and exon boundary consistency. In such approaches, exon identity is often determined through boundary matching. However, boundary-based matching is particularly sensitive to alignment noise, as small shifts at exon edges can create artificial junctions or disrupt junction consistency. In contrast, SCOTCH adopts an exon-centric strategy based on read–subexon alignment percentage (**Figure S13a**). For each subexon, the proportion of the exon segment aligned by a read is evaluated using two user-adjustable thresholds: alignment below *match_low* (10% as default) is considered unmapped, alignment above *match_high* (60% as default) is considered mapped, and intermediate cases are labeled as uncertain. This segment-based, percentage-driven approach is more robust to minor boundary shifts, alignment jitter, and local sequencing errors than discrete splice junction matching.

Handling noise during mapping typically requires thresholding. Most methods apply static, hard thresholds to determine whether a read supports an exon or junction. However, this introduces a fundamental trade-off: if the threshold is too strict, many reads fail to map to isoform annotations, artificially inflating the number of candidate novel isoforms or unmappable reads; if the threshold is too permissive, reads may become compatible with multiple isoforms or be incorrectly absorbed into existing annotations, thereby obscuring genuine novel isoform signals. Moreover, the optimal cutoff is gene- and data-dependent, especially in genes with complex isoform structures and fragmented sub-exons. A particular challenge arises from very small sub-exons (e.g., 1–20 bp), which frequently appear when annotations are decomposed into non-overlapping segments. For such short regions, mapping decisions are inherently unstable. If small exons are strictly enforced, minor alignment noise can create artificial conflicts and cause true read–isoform matches to be incorrectly rejected. Conversely, if small exons are ignored, the effective structural constraints are weakened, increasing the likelihood that a read becomes compatible with multiple isoforms and resulting in ambiguous or multi-mapping assignments. To address this, SCOTCH employs a dynamic thresholding strategy for small exon filtering (**Figure S13a**). Instead of using a fixed cutoff, the effective small-exon threshold is progressively increased from *small_exon_threshold_low* (0bp as default) to a maximum of *small_exon_threshold_high* (default as 80 bp). At each step, sub-exons shorter than the current threshold are masked and read–isoform compatibility is re-evaluated in a read-specific, strict-to-relaxed manner. The procedure stops once a compatible assignment is obtained, typically yielding a unique mapping. If no assignment is found even at the maximum threshold, the read is retained as confidently unmapped and forwarded for novel isoform reconstruction, reducing the risk that arbitrary cutoff choices generate spurious novel isoforms. This adaptive strategy improves mapping stability while preserving true structural variation.

**Comparison between exon-centric and splice-graph–based approaches for read-isoform mapping**

In this section, we compare exon-centric (e.g. SCOTCH) and splice-graph–based strategies for read–isoform mapping under a fixed isoform annotation. Most existing approaches represent isoform annotations as a splice graph. For example, in IsoQuant, splice junctions (introns) and transcript starting/end sites are treated as nodes, while exons serve as edges connecting these nodes. Read–isoform assignment is formulated as identifying a path in the graph whose junction sequence matches the observed splice junctions in the read. As illustrated in **Figure S13b**, Isoform1 (blue) corresponds to a path starting from terminal node T1 through junctions S1, S3, and S4 to T3, excluding nodes S2, S5, and T2. A read is assigned to Isoform1 if its splice junction pattern matches this path. On the other hand, exon-centric approaches decompose the annotation into non-overlapping sub-exons. Any isoform, known or novel, can be represented as a combination of sub-exons. Under this representation, Isoform1 corresponds to inclusion of E1, E2, E3, E4, E6, and E7, with E5 excluded. Read–isoform mapping is performed by evaluating compatibility between the read's sub-exon profile and the isoform's sub-exon combination.

Assuming the constructed splice graph and the derived sub-exon annotation are both accurate, the two representations are equivalent in ideal conditions. As shown in **Figure S13c**, reads originating from Isoform2 (green) with minimal noise will be correctly assigned by both splice-graph-based and exon-based approaches. However, differences emerge when there exist alignment artifacts. When local alignment artifacts occur near a splice boundary, such as those caused by sequence homology or clustered indels, exon elongation can occur, and two consecutive such artifacts may produce an apparent novel splice junction (S2') as shown in **Figure S13d**. In splice-graph–based mapping, only if both artifact segments (seg1 and seg2) fall below the algorithm's hard threshold can S2' be collapsed back to the annotated junction S2 for correct mapping. In the exon-centric framework, the segment (seg1) outside annotated exon space does not influence compatibility, and the impact of seg2 depends on the length of E3. If E3 is short, it may be masked during dynamic small-exon filtering, in which case seg2 does not affect compatibility. If E3 is long, seg2 must exceed the *match_high* threshold (default 60% alignment of E3 length) to be considered mapped. Because alignment artifacts rarely achieve substantial alignment over a relatively long exon, correct assignment is typically preserved.

Misalignment may also introduce short unannotated exon either within an intron region (**Figure S13e**, seg2) or at transcript ends (**Figure S13f**, seg1), sometimes co-occur with other alignment artifacts. In case 3 (**Figure S13e**), a false aligned exon (seg2) appears together with a junction drift (seg1), producing two apparent novel splice junctions (S1' and S2'). Under splice-graph–based mapping, correct assignment is recovered only if both artifacts fall below the hard threshold. As shown in case 4 (**Figure S13f**), false aligned exon (seg1) occurs in combination with read truncation at the transcript end, again creating an apparent novel splice junction (S0) with terminal node (T1) being missed, resulting in false incompatibility with the annotated isoform. In contrast, within the exon-centric framework, erroneous assignment requires stronger evidence: seg2 must exceed the *match_high* threshold relative to E2 in case 3 (**Figure S13e**), and seg1 must accumulate sufficiently read support to be considered a novel exon in case 4 (**Figure S13f**). Similarly, in case 5 (**Figure S13g**), multiple artifact types co-occur, including exon skipping and exon boundary shifts. In splice-graph–based approaches, any one of these segments exceeding the hard threshold may introduce apparent novel splice junctions or altered terminal nodes, leading to incorrect isoform rejection. In contrast, under the exon-centric framework, misassignment occurs only if these segments align sufficiently to annotated exon regions (e.g., exceeding the *match_high* threshold), making the error condition more stringent than in graph-based mapping.

As isoform complexity increases, multiple artifacts are more likely to co-occur within a single read. In splice-graph–based frameworks, each perturbation can introduce a new splice junction or alter the graph path, directly modifying the structural representation of the transcript. Consequently, combined artifacts are readily translated into additional junctions or path incompatibilities. In contrast, the exon-centric strategy simplifies isoform construction to evaluating read–subexon compatibility. Artifact segments occurring in annotated intronic regions are naturally excluded, and the remaining segments must provide sufficient exon-level coverage to alter the assignment. Because structural changes are not introduced unless exon-level support exceeds predefined thresholds, the error condition is inherently more stringent, reducing the likelihood that combined artifacts are escalated into novel transcript structures.

**Novel isoform reconstruction via iterative exon profile clustering**

Splice-graph–based methods typically begin novel isoform reconstruction by identifying splice junction coordinates from read alignments. These junctions, often clustered to account for minor positional variability, form the nodes of a splice graph from which transcript paths are inferred. As a result, structural inference is driven directly by splice-site detection: once a junction is accepted, it becomes an explicit structural element of the graph. Although graph simplification can merge similar junctions, variability at splice boundaries may either increase graph complexity (if retained) or cause reads carrying combined artifacts to become incompatible with simplified paths (if merged). Thus, structural uncertainty arises at the level of junction identification.

In contrast, SCOTCH begins by identifying exon segments rather than splice junctions. In enhanced mode, existing annotations are refined using read coverage information: novel exonic regions may be introduced in intronic space when sufficiently supported by reads, and annotated exons may be subdivided into non-overlapping exons according to observed coverage patterns (**Figure S13a**). This exon refinement step defines the segmental space in which isoforms are represented as combinations of sub-exons. Importantly, over-segmentation at this stage (e.g., identification of small or spurious sub-exons due to local noise) does not immediately create novel transcript structures. Instead, these segments serve as candidate building blocks whose relevance is evaluated in downstream clustering. Novel isoform reconstruction is then performed in exon-pattern space (**Figure 1c**). The input is a read–sub-exon compatibility matrix (−1, 0, 1), capturing exon inclusion patterns per read. Because reads originating from the same isoform may differ due to truncation or local mapping artifacts, SCOTCH first performs read clustering based on similarity of their exon-mapping profiles. This clustering step is intentionally permissive, allowing minor inconsistencies to be absorbed within clusters. For each sufficiently large cluster (e.g., >10 reads), a candidate novel isoform is inferred by majority voting across exon positions, producing a consensus exon combination. Reads are subsequently remapped to the updated isoform set, and unmapped reads are iteratively re-clustered and re-annotated. This loose-to-iterative process may initially generate multiple candidate isoforms, including redundant or partially truncated variants. A final global remapping step then evaluates all reads against the full candidate set. Because compatibility is assessed with dynamic exon-level thresholding, reads preferentially map to isoforms that better explain their exon patterns. Redundant annotations that lack stable read support are progressively eliminated.

Conceptually, although splice-graph–based approaches can absorb minor alignment variability during junction clustering, junctions associated with non-negligible artifacts remain distinct and are incorporated as separate nodes in the splice graph. If such junctions are retained, they increase graph complexity and may directly generate erroneous transcript annotations by introducing spurious splice sites and alternative paths. If they are aggressively merged or removed during graph simplification, reads carrying these perturbations may become incompatible with existing paths and remain unmapped. Importantly, junction identification and graph construction occur at an early stage and largely determine the structural search space. Errors introduced at this step, either by retaining artifact-derived junctions or by over-simplifying true variability, cannot be iteratively corrected during downstream read–isoform mapping. Consequently, structural misannotation may originate at the junction-identification stage and propagate throughout subsequent transcript inference. In contrast, SCOTCH performs exon segmentation and constructs transcript models in exon-pattern space through cluster-level consensus and iterative refinement. Candidate isoforms are not fixed at the moment of exon segmentation but are promoted only after consistent multi-exon support emerges within a read cluster and survives global remapping competition. Consequently, early over-segmentation or local noise at the exon-identification stage can be absorbed or filtered during clustering and refinement. Structural inference is therefore deferred until sufficient group-level evidence accumulates, effectively increasing the confidence threshold required for novel isoform annotation.

##### Evaluating novel isoform annotation in simulation study

We used gffcompare to compare GTF files of novel isoforms generated by each tool with the ground truth, establishing mappings between predicted and true novel isoforms within the same gene. To quantitatively assess annotation accuracy, we calculated precision and recall for each predicted–true isoform pair based on coordinate overlaps of isoform exons. For a predicted isoform p and true isoform t, we define:

$$Precision_{p,t} = \frac{Length\ of\ overlapping\ exon\ regions\ between\ p\ and\ t}{Total\ exon\ length\ of\ p}$$

$$Recall_{p,t} = \frac{Length\ of\ overlapping\ exon\ regions\ between\ p\ and\ t}{Total\ exon\ length\ of\ t}$$

$$F1_{p,t} = \frac{2\, Precision_{p,t}\, Recall_{p,t}}{Precision_{p,t} + Recall_{p,t}}$$

Isoforms with F1 scores exceeding 0.8 were considered correctly annotated. We defined precision in a prediction-centric manner as the proportion of predicted novel isoforms correctly matched to any ground-truth novel isoform:

$$Precision = \frac{\#\,(predicted\ novel\ isoforms\ with\ F1_{p,t} \geq 0.8)}{\#\,(predicted\ novel\ isoforms)}$$

Similarly, recall was defined in a truth-centric manner as the proportion of ground-truth novel isoforms successfully recovered by at least one predicted isoform:

$$Recall = \frac{\#\,(ground - truth\ novel\ isoforms\ with\ \max F1_{p,t} \geq 0.8)}{\#\,(ground - truth\ novel\ isoforms)}$$

##### Reference


1. Mikheenko, A., Prjibelski, A.D., Joglekar, A. & Tilgner, H.U. Sequencing of individual barcoded cDNAs using Pacific Biosciences and Oxford Nanopore Technologies reveals platform-specific error patterns. *Genome Res* **32**, 726–737 (2022).
